# Rapid Assessment of the Temporal Function and Phenotypic Reversibility of Neurodevelopmental Disorder Risk Genes in *C. elegans*

**DOI:** 10.1101/2021.10.21.465355

**Authors:** Lexis D. Kepler, Troy A. McDiarmid, Catharine H. Rankin

## Abstract

Hundreds of genes have been implicated in neurodevelopmental disorders. Previous studies have indicated that some phenotypes caused by decreased developmental function of select risk genes can be reversed by restoring gene function in adulthood. However, very few risk genes have been assessed for adult reversibility. We developed a strategy to rapidly assess the temporal requirements and phenotypic reversibility of neurodevelopmental disorder risk gene orthologs using a conditional protein degradation system and machine vision phenotypic profiling in *Caenorhabditis elegans*. Using this approach, we measured the effects of degrading and re-expressing orthologs of 3 neurodevelopmental risk genes *EBF3, BRN3A*, and *DYNC1H1* across 30 morphological, locomotor, sensory, and learning phenotypes at multiple timepoints throughout development. We found some degree of phenotypic reversibility was possible for each gene studied. However, the temporal requirements of gene function and degree of phenotypic reversibility varied by gene and phenotype. The data reflects the dynamic nature of gene function and the importance of using multiple time windows of degradation and re-expression to understand the many roles a gene can play over developmental time. This work also demonstrates a strategy of using a high-throughput model system to investigate temporal requirements of gene function across a large number of phenotypes to rapidly prioritize neurodevelopmental disorder genes for re-expression studies in other organisms.

**SUMMARY STATEMENT:** We developed a strategy that combines a conditional and reversible protein degradation system with our high-throughput machine vision tracking system to assess the temporal windows of gene function and reversibility of phenotypic disruptions associated with neurodevelopmental disorder risk gene orthologs using *C. elegans*. Using this approach, we assessed 3 genes (*unc- 3*, *unc-86*, and *dhc-1)* and found that post-embryonic rescue was possible for each gene and each phenotypic feature class assessed. Re-activation of certain genes was able to reverse multiple phenotypic disruptions late into development without inducing novel phenotypes, prioritizing them for further study.

## INTRODUCTION

Neurodevelopmental disorders such as Autism Spectrum Disorder (ASD) and Intellectual Disability (ID) are highly genetically heterogeneous and are accompanied by a range of cognitive and behavioural phenotypes including sensory processing and learning impairments (American Psychiatric Association, 2013; Boyle et al., 2011; de la Torre-Ubieta et al., 2016; Iakoucheva et al., 2019; Sanders et al., 2019). More severe cases of these disorders can cause significant challenges for affected individuals and their families (Boyle et al., 2011; de la Torre-Ubieta et al., 2016; Iakoucheva et al., 2019; Sanders et al., 2019). Recently, there has been remarkable progress in identifying genetic risk factors that contribute to diverse neurodevelopmental disorders, with hundreds of genes now implicated in ASD and ID alone (de la Torre-Ubieta et al., 2016; De Rubeis et al., 2014; Deciphering Developmental Disorders Study, 2015; Iakoucheva et al., 2019; Iossifov et al., 2014; Sanders et al., 2019, 2015; Satterstrom et al., 2020; Vissers et al., 2016). Variants in the majority of these genes (e.g. 89/102 ASD-associated genes; 87%) are thought to confer risk through haploinsufficiency as the individual carries one loss-of-function allele with insufficient residual function from the remaining copy (Satterstrom et al., 2020). The identification of how these variants contribute to disorder pathology suggests re-expression therapies, where a second functional allele is introduced to restore protein levels to compensate for the decreased function of the faulty allele, could be a viable treatment option.

Historically, it was assumed that any treatment targeting neurodevelopmental disorder risk genes would need to be administered very early in development to be effective. This long-held assumption was challenged by reports that re-expression of several risk gene orthologs could reverse multiple altered neurophysiological and/or behavioural phenotypes in adult mice (Creson et al., 2019; Ehninger et al., 2008; Gao et al., 2020; Guy et al., 2007; Mei et al., 2016; Vogel-Ciernia et al., 2013; Zeier et al., 2009). In addition, inactivating orthologs of some of these risk genes in adult mice could also induce the phenotypic impairments previously associated only with altered gene function during development (Ehninger et al., 2008; Guy et al., 2007). Together, these findings suggest there may be a degree of temporal flexibility in the neurodevelopmental processes these gene contribute to, and that some genes typically associated with neurodevelopment may continue to have important functions well into adulthood.

The handful of reports that show the possibility of phenotypic reversibility with gene reactivation in adult mice offer critical insights into which genes are promising candidates for future re-expression-based therapies. However, because of limitations including cost, developmental rate, and technical difficulties (e.g. injection of viral vectors for large numbers of animals, etc.) very few risk genes have been tested for adult phenotypic reversibility. Further, the rapidly growing number of risk genes identified in recent years has exacerbated this problem and drastically increased the need for candidate prioritization to better direct research efforts. While most neurodevelopmental disorder risk genes are highly expressed early in pre-natal development (Jin et al., 2020; Parikshak et al., 2013; Satterstrom et al., 2020; Willsey et al., 2013), many continue to be expressed well into adulthood, and we currently do not know if the relationship between temporal expression patterns and inferred temporal functional windows are a significant predictor of whether a gene will be suitable for re-expression therapies. Assessing the phenotypic reversibility of neurodevelopmental disorder risk genes in more high-throughput model organisms offers the ability to rapidly screen a large number of genes to aid in prioritizing risk genes for further study in mammalian models.

The nematode *Caenorhabditis elegans* offers multiple advantages to systematically assess the temporal requirements and phenotypic reversibility of neurodevelopmental disorder risk genes. *C. elegans* have orthologs for a high number of neurodevelopmental risk genes (e.g. >80% of high-confidence ASD risk genes (McDiarmid et al., 2020)) and these genes have repeatedly been shown to be so well-conserved that in many cases expression of the human risk gene can compensate for loss of the *C. elegans* ortholog (Kaletta and Hengartner, 2006; Levitan et al., 1996; McDiarmid et al., 2018; Post et al., 2020). *C. elegans* have rapid development, growing from egg through well-characterized larval stages (L1, L2, L3, and L4) to egg-laying adults within 3 days. The hermaphroditic reproduction of *C. elegans* enable large colonies of genetically identical animals to be rapidly and cheaply cultivated. In addition, multiple genomic tools are available to precisely control the spatial and/or temporal activity of genes *in vivo* (Ashley et al., 2021; Au et al., 2019; Dickinson and Goldstein, 2016; Nance and Frøkjær-Jensen, 2019; Wang et al., 2017; Zhang et al., 2015). Lastly, the phenotypic profiles of hundreds of animals can be simultaneously assessed using automated tracking systems which capture and analyze the impact of genetic perturbations on morphological, sensory, and learning behaviours in real time (Husson, Steuer Costa, Schmitt, and Gottschalk, 2012; McDiarmid et al., 2018; Swierczek et al., 2011).

A behaviour that has been increasingly used in high-throughput model organisms to investigate the biological function of neurodevelopmental disorder risk genes and functional impact of disorder-associated variants is habituation (Kepler et al., 2020). Habituation is a highly conserved form of non-associative learning observed as a decrement in responding to a repetitive stimulus (Rankin et al., 2009; Thompson and Spencer, 1966). Habituation is thought to be an important building block for higher cognitive functions and enable ongoing shifts in behavioral strategy (McDiarmid et al., 2019; Schmid et al., 2015). Alterations in the ability to habituate have been reported in ASD, ID, and Schizophrenia (McDiarmid et al., 2017) and are hypothesized to contribute to more complex behavioural symptoms (Green et al., 2015; Kavšek, 2004; Kleinhans et al., 2009; Massa and O’Desky, 2012; Williams et al., 2013).

Here, we developed a strategy to assess the phenotypic reversibility and temporal functional windows of three neurodevelopmental disorder risk genes *in vivo* using CRISPR-Cas9 Auxin Inducible degradation (AID) and high-throughput machine vision phenotyping (**Fig. 1**). We took advantage of the genetic tractability and rapid, stereotyped development of *C. elegans* to precisely investigate the effects of degrading and re-expressing the proteins of interest at multiple developmental time points in thousands of age-synchronized genetically identical animals. We found that some level of phenotypic reversibility was possible for each risk gene if re-expression occurred early in post-embryonic development, but only re-expressing *EBF3•unc-3* and *DYNC1H1•dhc-1* could reverse multiple phenotypic alterations later in life. More broadly, we provide an adaptable strategy and important examples/criteria that illustrate a path towards prioritizing neurodevelopmental disorder risk genes for further study and therapeutic development.

**Figure 1.**
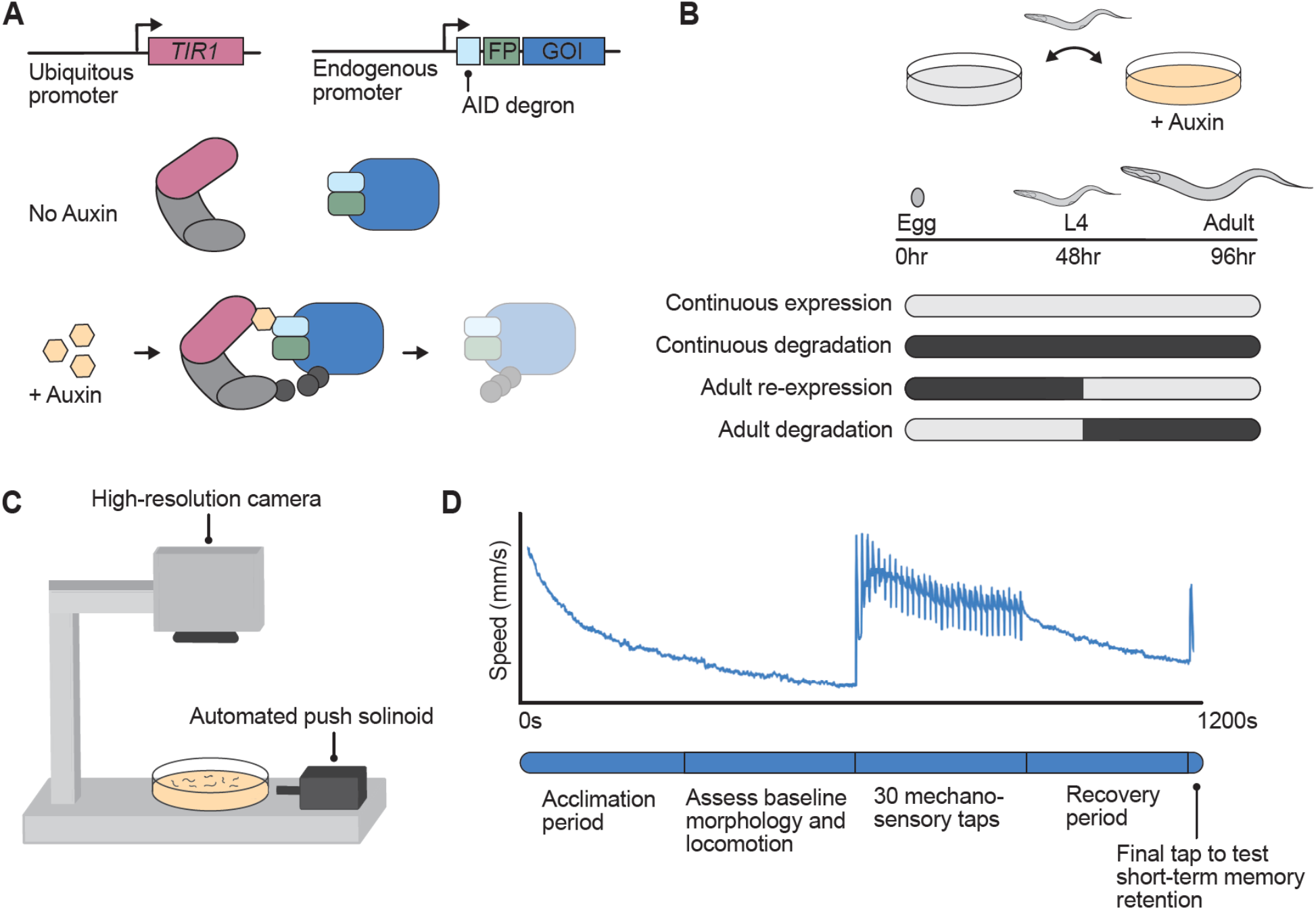
A pipeline to assess the temporal requirements and phenotypic reversibility of neurodevelopmental disorder risk gene orthologs using the AID system and machine vision phenotypic profiling in Caenorhabditis elegans. **A)** The Auxin-Inducible Degradation (AID) system is a powerful approach that enables temporal and spatial control of protein depletion. CRISPR-Cas9 is used to tag the gene of interest (GOI) with the AID degron along with a fluorescent protein (FP) to visualize protein expression *in vivo*. In the presence of the small molecule Auxin, TIR1 (an E3 ubiquitin ligase) associates with the AID degron, recruiting endogenous proteosomes to degrade the ubiquitinated protein of interest. **B)** Temporal degradation conditions were created by manually transferring animals on and off Petri plates containing Auxin to inactive or restore gene function at specific timepoints in development or adulthood. **C)** The effects of protein degradation and re-expression across 26 morphological, locomotor, and sensory and learning phenotypes were objectively quantified in hundreds of animals simultaneously using a machine vision tracking system throughout a short-term mechanosensory habituation paradigm (D).

## RESULTS

We selected three neurodevelopmental disorder risk genes (as identified by Simons Foundation Autism Research Initiative (Abrahams et al., 2013), Satterstorm et al. (2020), and/or literature search) for reversibility analysis based on the availability of *C. elegans* strains that contain a neurodevelopmental disorder risk gene ortholog tagged with the Auxin-Inducible Degron at the endogenous locus using CRISPR-Cas9 (see methods). The AID system relies on tagging the gene of interest with a short degron peptide tag as well as transgenic expression of *TIR1* which is an E3 ubiquitin ligase typically found only in plants (Nishimura et al., 2009; Zhang et al., 2015). In the presence of the plant hormone Auxin, *TIR1* can associate with the AID degron, adding a poly-ubiquitin chain to the protein of interest causing it to be degraded by the proteasome (Nishimura et al., 2009; Zhang et al., 2015) (**Fig. 1A**). We chose to use the AID system as it enables temporal control of protein degradation, which can be reversed by transferring populations of worms to culture plates without Auxin (McDiarmid et al., 2020; Zhang et al., 2015). Since the AID degron is tagged to the endogenous loci, protein expression is restored using the native regulatory machinery, therefore bypassing the biological confounds associated with conventional approaches that rely on overexpression. Importantly, our lab and others have shown that Auxin exposure does not cause any overt effects on *C. elegans* morphology, locomotion, short-term learning, or mechanosensory processing phenotypes (McDiarmid et al., 2020; Zhang et al., 2015).

We systematically assessed the functional consequence of multiple developmental degradation time windows *in vivo* by transferring animals on or off plates containing Auxin at precise time points in *C. elegans* development (**Fig. 1B**). We used our high-throughput machine vision tracking system, the Multi-Worm Tracker (MWT) (Swierczek et al., 2011), to quantify 26 phenotypes spanning morphology, baseline locomotion, sensory responding, and learning while animals were subjected to a short-term mechanosensory habituation behavioral paradigm (**Fig. 1C & D**). Our phenotypic features included multiple measures of mechanosensory responding and habituation learning, as both are disrupted across neurodevelopmental disorders (Green et al., 2019, 2015; McDiarmid et al., 2017; Williams et al., 2013), and because we have previously shown that different components of a single behavioral response can be mediated by genetically dissociable underlying mechanisms (Ardiel et al., 2018; Kindt et al., 2007; McDiarmid et al., 2019; Randlett et al., 2019). Inclusion of a range of phenotypes not only aids in characterizing gene function across development, but also enables any unexpected phenotypes caused by protein re-expression to be captured.

### *The transcription factor EBF3●unc-3* displays a reciprocal pattern of phenotypic induction and reversibility across development

The first neurodevelopmental disorder risk gene we assessed was *EBF3•unc-3,* which is a highly conserved transcription factor. In *C. elegans*, *unc-3* that acts to specify the identity of different neuronal classes by initiating and maintaining the expression of class-specific effector genes (**Fig. 2A**) (Kratsios et al., 2017; Prasad et al., 2008, 1998). Variants in *EBF3* have been associated with multiple neurodevelopmental disorders including ASD and ID, and are thought to confer risk through haploinsufficiency or by interfering with DNA binding (Chao et al., 2017; Lopes et al., 2017; Sleven et al., 2017; Tanaka et al., 2017). In *C. elegans*, *unc-3* loss-of-function results in severe locomotion and coordination defects caused by undifferentiated/abnormal identity of cholinergic motor neurons in the ventral nerve cord (Brenner, 1974; Feng et al., 2020; Kratsios et al., 2017; Prasad et al., 1998).

**Figure 2.**
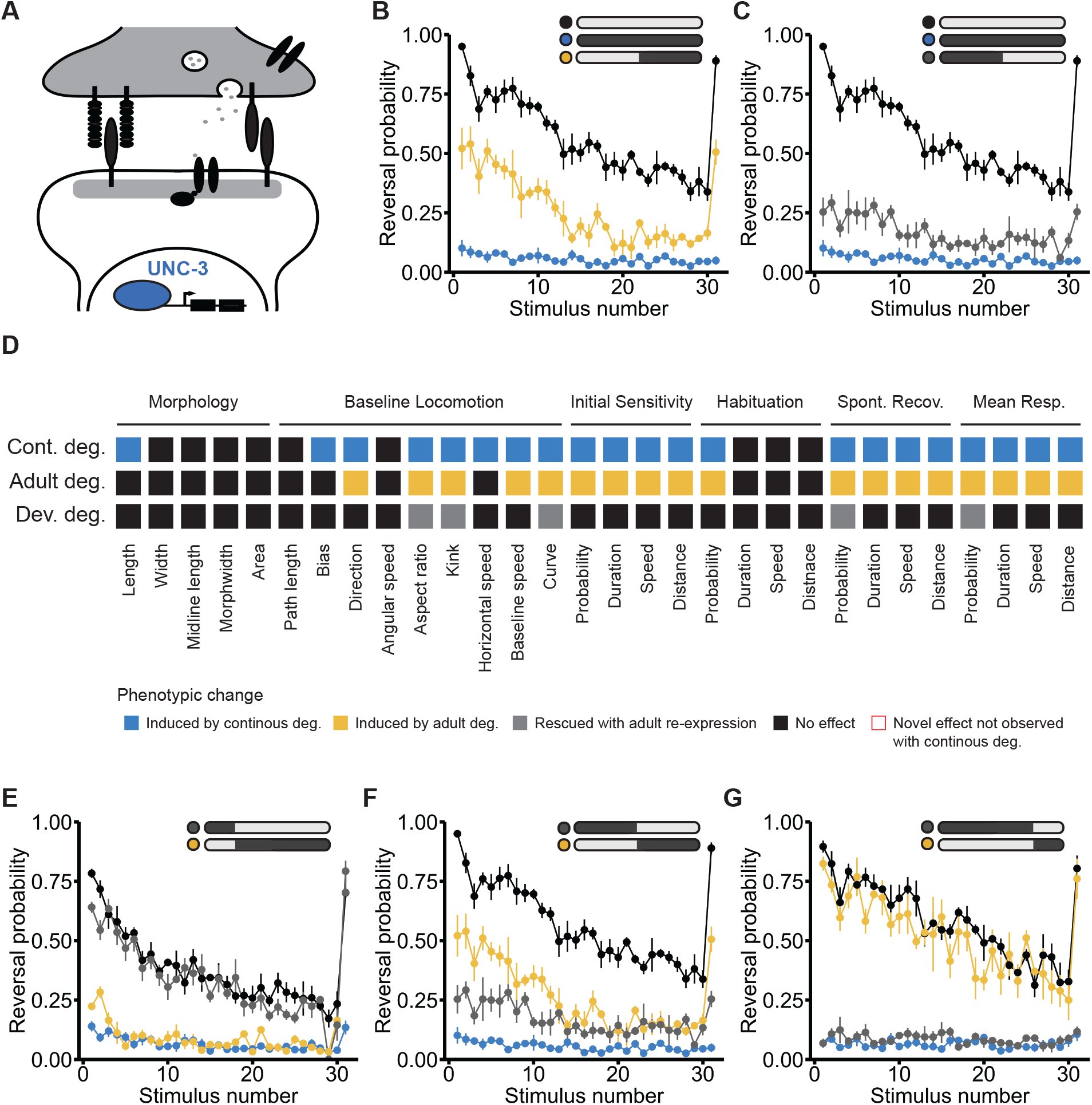
The transcription factor EBF3●unc-3 displays a reciprocal pattern of phenotypic induction and reversibility across development. A) The transcription factor *EBF3•unc-3* acts to specify neuronal identity. B) Continuous degradation *unc-3* (blue) impaired the animals’ ability to respond to mechanosensory stimuli compared to the no-Auxin control (black). Starting Auxin exposure at L4 partially induced this impairment. C) Ending Auxin exposure after L4 (48 hrs post- hatch) partially rescued the impairment in response probability. D) Full phenotypic profile of *unc- 3*, indicating all phenotypes induced by continuous degradation (blue), induced by adult-specific degradation (starting at L4, yellow), and rescued with adult re-expression (gray). E) Ending Auxin exposure at L2 (24 hrs post-hatch, gray) almost completely rescued the impairment, F-G) with the level of phenotypic rescue decreasing with later onset of UNC-3 re-expression. E) Starting Auxin exposure at L2 (yellow) induced an impairment level similar to the continuous degradation group, F-G) with the degree of phenotypic impairment decreasing with later onset of UNC-3 degradation.

In our paradigm, continuous degradation of UNC-3 from egg through to adulthood produced uncoordinated locomotion, an inability to respond to mechanosensory stimuli, and severe alterations in several other morphological and behavioral phenotypes (**Fig. 2B & D and Fig 3**). Early adult inactivation of *unc-3* (achieved by beginning Auxin exposure at the final larval stage, L4) induced impairments for the majority of affected phenotypes, supporting previous findings that *unc-3* function is continuously required throughout development (Feng et al., 2020; Kratsios et al., 2017; Li et al., 2020) (**Fig. 2B**). Importantly, impairments in mechanosensory response probability and several other altered phenotypes were partially rescued in animals when UNC-3 was degraded during development but was re-expressed from the endogenous locus starting in early adulthood (animals taken off Auxin immediately after L4) (**Fig. 2C**).

**Figure 3.**
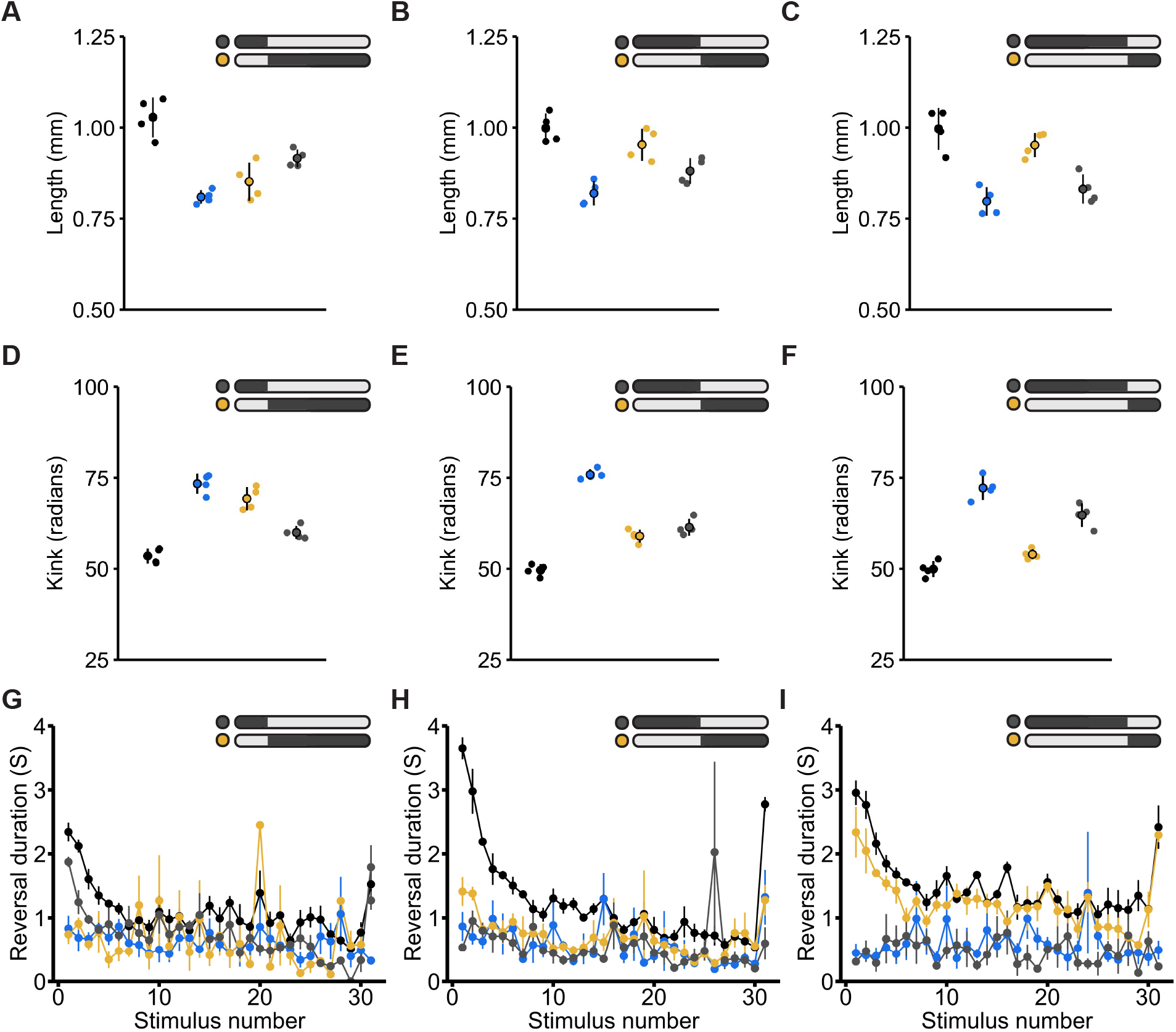
EBF3•unc-3 shows diverse temporal patterns of phenotypic reversibility across morphological, locomotor, and mechanosensory response phenotypes. A) The no-Auxin control group is depicted in black and continuous degradation group is depicted in blue for all panels. Altered animal length could be partially rescued with early post embryonic re-expression (starting at L2/24 hrs post-hatch) or fully induced with early post embryonic degradation. B) The degree of rescue and impairment of animal length increasingly diminished if UNC-3 was re- expression or degraded at L4 (48 hrs post-hatch) C) or in adulthood (72 hrs post-hatch). D) Similarly, impairments in kink could be fully rescued with early post embryonic re-expression, E-F) but degree of rescue diminished with later re-expression. D) Degrading UNC-3 starting at L2 resulted in impaired kinkiness to a level similar to the continuous degradation control group, E-F) the degree of impairment lessened with later onset of degradation. G-I) Impairments in response duration could not be rescued with UNC-3 re-expression across any of the tested temporal conditions. G-H) Degrading UNC-3 in early post embryonic development or development (L4) induced impairments similar to the continuous degradation condition, I) yet duration impairments were not induced with UNC-3 degradation starting at 72 hrs post-hatch.

Our initial findings of partial rescue of some phenotypes following protein re-expression starting at early adulthood motivated us to explore whether earlier restoration of *unc-3* function would produce more effective rescue. Re-expression of UNC-3 during late post-embryonic development (ending Auxin exposure at L2) resulted in an almost complete rescue of impairments in response probability (**Fig. 2E**). Further, starting UNC-3 degradation during late post-embryonic development produced more severe impairments which were similar to the continuous degradation control, suggesting *unc-3* plays a critical role between L2 and L4 for reversal probability (**Fig. 2E**). Lastly, we explored the phenotypic consequences of exposing or removing 3-day old animals (young adult- 72 hrs post-hatch) from Auxin. Re-expression of UNC-3 in adulthood did not rescue impairments in response probability, reaffirming that the crucial functional period of *unc-3* occurs during development. In line with this, starting UNC-3 degradation at 72 hrs post-hatch did not induce impairments, suggesting that the role of *unc-3* in maintaining the expression of terminal identity genes throughout the lifespan may not be required for normal behaviour once the nervous system has fully developed.

Assessment across the three temporal conditions revealed a striking reciprocal pattern of *unc-3* temporal function for response probability (**Fig. 2E-G**). The degree of phenotypic impairment for response probability induced by inactivation of *unc-3* at a given developmental time point almost perfectly mirrored the degree of reversibility possible with re-expression (**Fig. 2E-G**). While some other phenotypes followed this reciprocal pattern others did not (**Fig. 3A-F**). For example, reversal duration seems to be mediated by a mechanism that acts earlier in development, with reversibility only possible with early re-expression (**Fig. 3G-I**). Taken together, these results clearly demonstrate that the temporal windows of gene function can drastically vary by phenotype, and that both the degree of reversibility and number of reversible phenotypes can be influenced by how early re-expression occurs in development. Importantly, these findings reveal that *unc-3* expression can rescue multiple phenotypic alterations relatively late in life, prioritizing this gene as a candidate for further study.

### *BRN3A●unc-86* specifically impairs mechanosensory response probability and displays a reversibility window restricted to early post-embryonic development

*BRN3A•unc-86* is a POU-type transcription factor that plays conserved roles in the initiation and maintenance neuronal identity across species (Badea et al., 2009; Serrano-Saiz et al., 2018; Xiang et al., 1995; Zou et al., 2012). Variants in *POUF4/BRN3A* have been associated with abnormal development of sensory structures, including auditory (Huang et al., 2001) and visual cells (Badea et al., 2009). In *C. elegans*, *unc-86* is thought to be required across the lifespan to maintain the expression of terminal identity genes in multiple neuronal subtypes (Serrano-Saiz et al., 2018; Sze and Ruvkun, 2003) (**Fig. 4A**).

**Figure 4.**
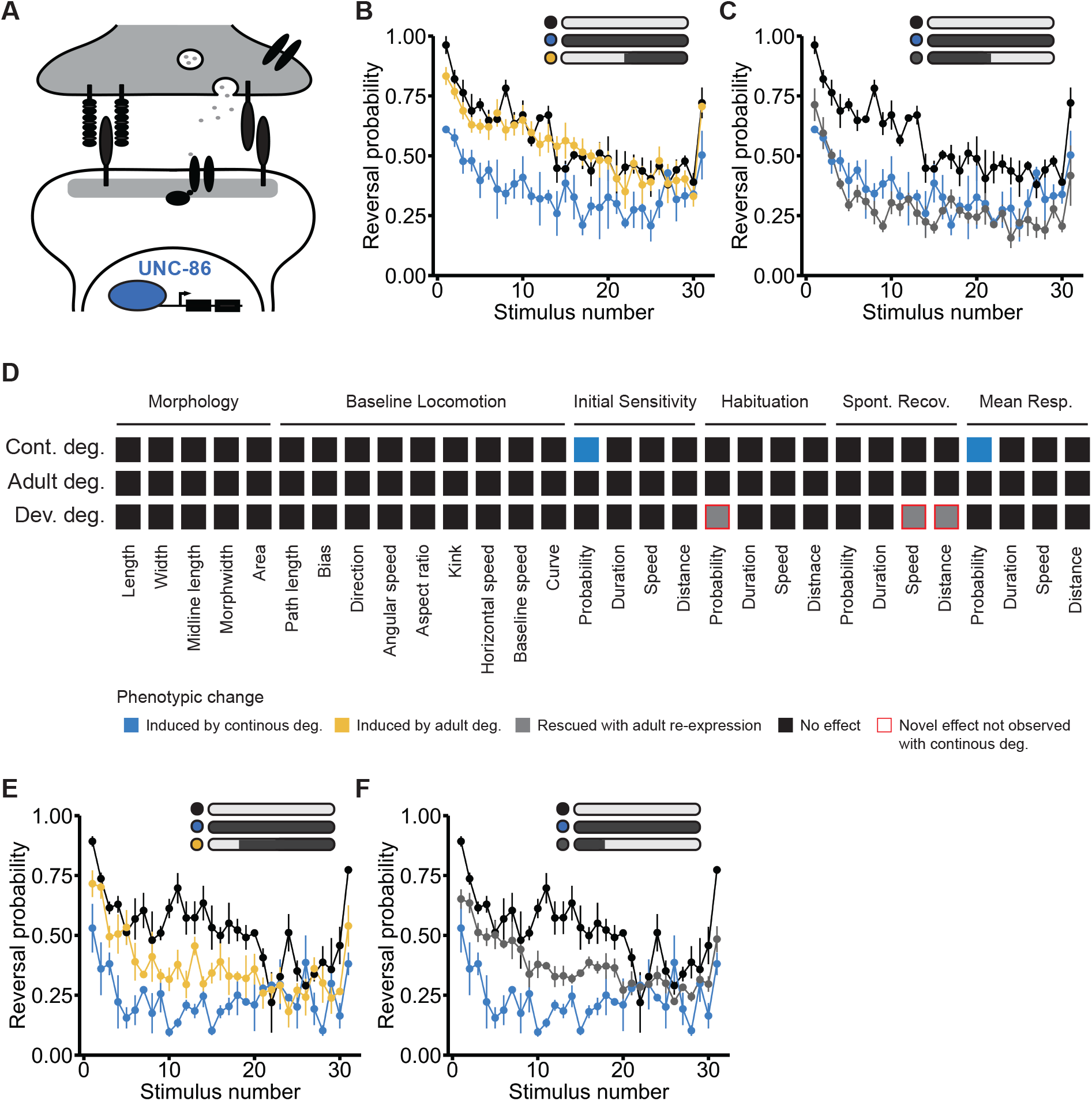
BRN3A●unc-86 specifically impairs mechanosensory response probability and displays a reversibility window restricted to early post-embryonic development. A) The transcription factor *BRNA3•unc-86* acts maintain the expression of terminal identity genes in multiple neuron types. B) Continuous degradation of UNC-86 (blue) specifically impaired response probability to mechanosensory stimuli compared to animals that were not exposed to Auxin (black). Staring Auxin exposure at L4 (48 hrs post-hatch, yellow) did not significantly induce phenotypic impairments. C) Ending Auxin exposure at L4 (gray) did not rescue impairments in response probability. D) Full phenotypic profile of *unc-86*, indicating all phenotypes induced by continuous degradation (blue), induced by adult-specific degradation (starting at L4, yellow), and rescued with adult re-expression (gray). E) Exposing animals to Auxin starting at L2 (yellow) induces impairments in response probability. F) Ending Auxin exposure at L2 (24 hrs post-hatch, gray) enabled phenotypic rescue.

Analysis of 30 quantitative phenotypes revealed that continuous degradation of UNC-86 specifically impaired mechanosensory response probability (**Fig. 4B**). Animals were hyporesponsive to mechanosensory stimuli, with a lower likelihood of responding throughout the tracking session, while other parameters of the reversal response (e.g. duration and speed) were not affected. Re-expression of UNC-86 in early adulthood did not rescue these impairments (**Fig. 3C**) and exposing animals to Auxin from L4 onwards did not induce the hyporesponsive phenotype seen with continuous degradation (**Fig. 4B**). Together, these findings suggest a primarily early developmental role for *unc-86* in regulating mechanosensory responses. While we found that impairing *unc-86* from L4 onwards did not induce any significant impairments, a previous study found that inactivating *unc-86* at L4 using a temperature-sensitive allele resulted in impaired chemotaxis to multiple odorants (Sze and Ruvkun, 2003). These differences may suggest that *unc-86* has distinct temporal functional windows in different neuronal classes (i.e. *unc-86* is continuously required for the function of chemotaxis neurons but is only required in early development for function of mechanosensory neurons).

We next investigated the phenotypic consequences of degrading and re-expressing UNC- 86 starting at an earlier developmental time point (specifically 24 hours after age-synchronization during late post-embryonic development; beginning after L2). Restoration of *unc-86* expression beginning at this earlier time point was sufficient to completely rescue impairments in mechanosensory hyporesponsivity (**Fig. 4E**). In contrast, we found that degrading UNC-86 from this same time point onwards also impaired response probability (**Fig. 4F**). These results suggest the window of phenotypic reversibility for *unc-86* extends into early post-embryonic development. Moreover, these results indicate that as long as *unc-86* has played its role in neurodevelopment in this early critical window, it is no longer required for normal mechanosensory responding in adulthood.

### Ubiquitous degradation of the essential protein *DYNC1H1●dhc-1* in adult animals reveals specific roles in mechanosensory responding and habituation

*DYNC1H1•dhc-1* is an essential motor protein implicated in diverse processes including cell division and cargo transport along microtubules (e.g. retrograde axonal transport in neurons) (Cianfrocco et al., 2015) (**Fig. 5A**). Variants in *DYNC1H1* have been implicated in several neurodevelopmental disorders including ID and ASD (Satterstrom et al., 2020; Willemsen et al., 2013). Determining the biological functions of *DYNC1H1•dhc-1* throughout development has been challenging as dynein loss-of-function results in early embryonic lethality in multiple model organisms (Hamill et al., 2002; Harada et al., 1998; Howell and Rose, 1990; Mains et al., 1990; Robinson et al., 1999). As a result, the role of *DYNC1H1•dhc-1* in behaviour remains relatively uncharacterized.

**Figure 5.**
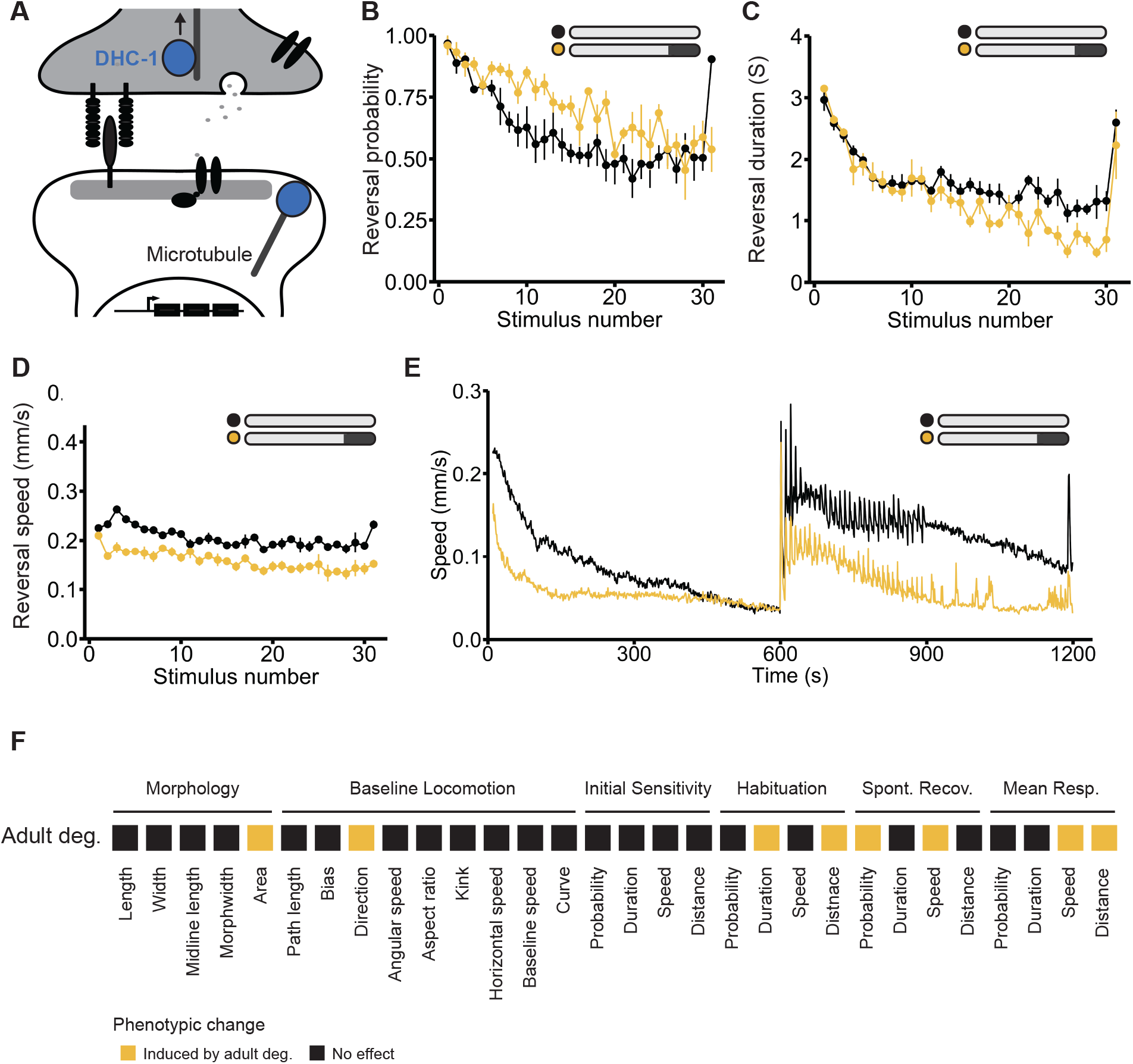
Ubiquitous degradation of the essential protein DYNC1H1●dhc-1 in adult animals reveals specific roles in mechanosensory responding and habituation. A) The essential gene *DYNC1H1•dhc-1* acts in cargo transport and stabilization of microtubule dynamics. B) Starting Auxin exposure in early adulthood (72 hrs post-hatch) did not impair response probability but C) did deepen habituation of response duration and D) decreased response speed compared to the no-Auxin control animals (black). E) Degrading dynein in adult animals (yellow) decreased average speed during the acclimation period and caused deeper habituation of response speed across the mechanosensory stimuli resulting in a lower average speed during the rest period post mechanosensory stimulation. F) Full phenotypic profiles of *dhc-1*, indicating all phenotypes induced by adult-specific degradation starting at 3 days post synchronization (yellow).

Here, we used the AID system to ubiquitously degrade DHC-1. As expected from loss-of-function alleles, continuous degradation of dynein was lethal. Degrading dynein in early adulthood (starting Auxin exposure immediately after L4) also resulted in lethality, indicating that dynein function remains essential throughout the late stages of *C. elegans* development. To determine whether dynein function is essential in adulthood, we ubiquitously degraded dynein in 3-day old adults (beginning 72 hrs post-hatch). We found that degrading dynein in adulthood was not lethal, allowing us an opportunity to investigate the biological functions of dynein in adult animals.

Despite its broad expression and essential role in early development, degradation of dynein later in life revealed surprisingly specific roles in adult behaviour (**Fig. 5A-F**). Adult-specific degradation of dynein did not cause severe alterations in morphology or baseline locomotion. Instead, only select components of the mechanosensory reversal response were affected while others were left intact. Response probability was unaffected, as the proportion of worms that responded to each mechanosensory stimulus was similar to the no Auxin control (**Fig. 5B**). However, degrading dynein at 72 hrs post-hatch caused animals to display deeper habituation of response duration (**Fig. 5C**) and a slower response speed compared to animals not exposed to Auxin (**Fig. 5D**). Interestingly, the absolute speed trace across the entire experiment shows animals have the ability to respond as fast as control animals but decrement their response speed faster with repeated stimulation (**Fig. 5E**). These findings suggest that, in adulthood, dynein may function to promote normal habituation of response duration and speed, but not response probability. Taken together, these results support the hypothesis that different components of habituation can be mediated by distinct mechanisms, and reveal novel, adult-specific roles for *DYNC1H1•dhc-1* in mechanosensory responding and habituation (**Fig. 5F**)

### Pan-neuronal degradation of *DYNCH1●dhc-1* is not lethal and causes multiple habituation impairments with distinct reversibility profiles

To further investigate the role of dynein across development, we took advantage of the ability to activate the AID system cell specifically and obtained a line of *C. elegans* that allowed for specific and reversible degradation of dynein only in neurons by driving *TIR1* expression under the *rab-3* promoter (**Fig. 6A**). Continuous pan-neuronal degradation of dynein did not cause lethality, offering us an unprecedented opportunity to determine the phenotypic consequences of decreased dynein function in the nervous system throughout development and whether the resulting impairments were reversible.

**Figure 6.**
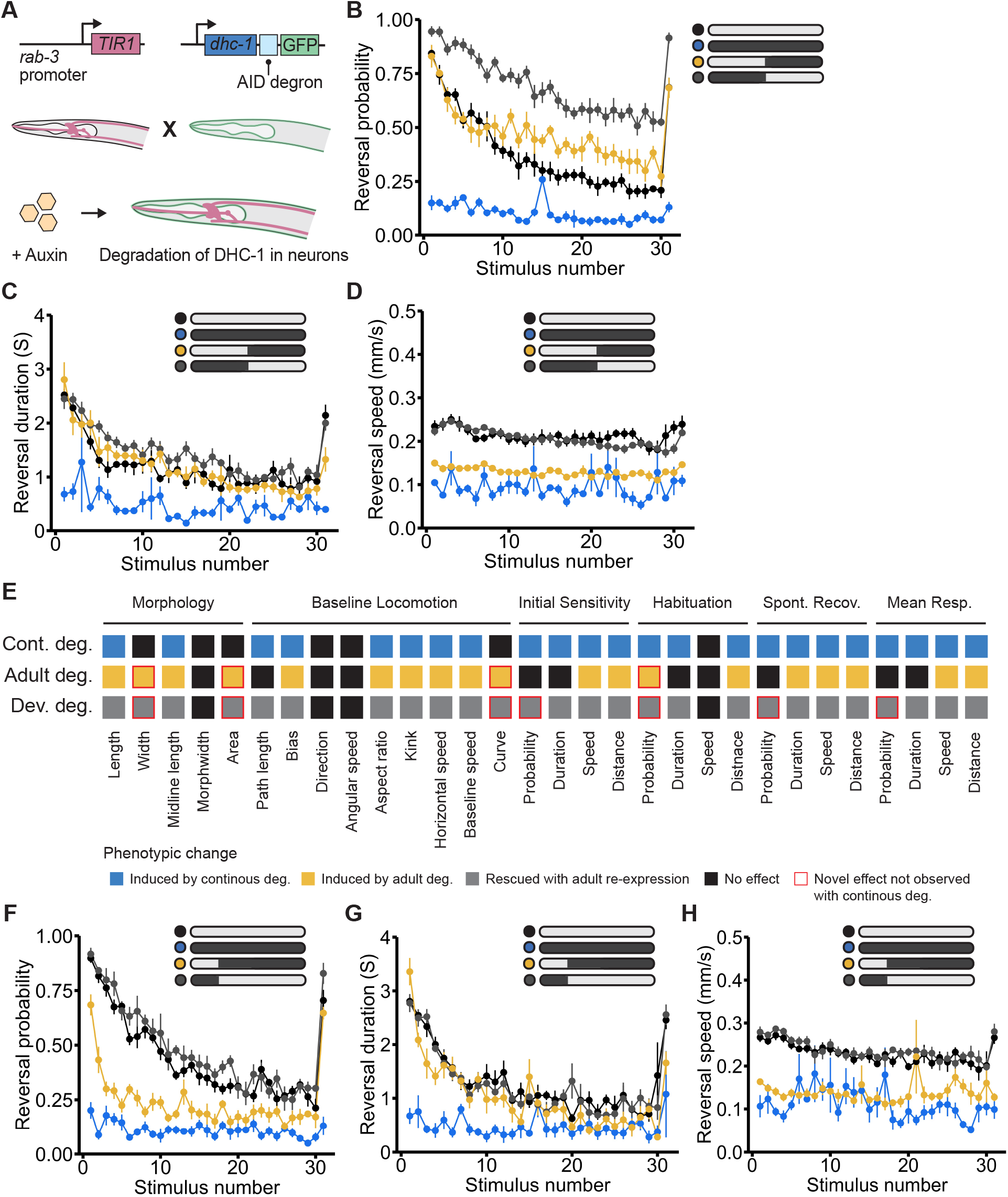
Pan-neuronal degradation of DYNCH1●dhc-1 is not lethal and causes multiple habituation impairments with distinct reversibility profiles. A) Pan neuronal degradation of dynein was achieved by crossing the *dhc-1(ie28[dhc-1::degron::GFP])* strain with a strain where *TIR1* expression is driven by the *rab-3* promoter. B) Continuous degradation of neuronal dynein (blue) impaired response probability compared to the no-Auxin control group (black). Re-expressing (gray) or degrading (yellow) neuronal dynein starting at L4 (48 hrs post-hatch) caused animals to be hyperresponsive to mechanosensory stimuli. C and D) Impairments in response duration and response speed could be rescued with re-expression of dynein starting at L4 (gray). Starting degradation of neuronal dynein at L4 (yellow) did not induce significant impairments in response duration but did induce impairments in response speed. E) Full phenotypic profile of *dhc-1*, indicating all phenotypes induced by continuous degradation (blue), induced by adult-specific degradation (starting at L4, yellow), and rescued with adult re-expression (gray). F) Exposing animals to Auxin at L2 (24 hours post-hatch) impaired response probability (yellow) and ending Auxin exposure at L2 rescued impairments in response probability (gray). G and H) Impairments in response duration and speed were rescued with re-expression of dynein starting at L2 (gray). Starting Auxin exposure at L2 (yellow) did not affect response duration, but did impair response speed.

Continuous pan-neuronal degradation of dynein caused a broad range of sensory responding impairments, including a low probability of responding to mechanosensory stimuli, rapid habituation of response duration, and lower response speed (**Fig. 6B-E and Fig. 7A-D**). In this case, multiple phenotypes showed distinct temporal functional windows and reversibility patterns. Re-expression of dynein in early adulthood (ending Auxin exposure immediately after L4) did not restore normal mechanosensory responding, but instead resulted in severely impaired habituation of response probability (animals were hyperresponsive and did not learn to decrease their likelihood of responding to repeated stimuli) (**Fig. 6B**). Adult pan-neuronal degradation of dynein (beginning Auxin exposure at L4) did not alter initial response probability but did decrease habituation of response probability (**Fig. 6B**). For response duration, adult re-expression of dynein was sufficient to fully rescue the impairment seen with continuous degradation, but adult degradation did not induce the response duration impairment (**Fig. 6C**). Consistent with our findings for ubiquitous dynein degradation in 72 hr old adult animals, continuous degradation of pan neuronal dynein also caused animals to exhibit slower reversal speed. The speed impairment could be rescued with dynein re-expression in early adulthood and was also induced by adult degradation (**Fig. 6D**) suggesting that dynein is continuously required in neurons to mediate response speed.

**Figure 7.**
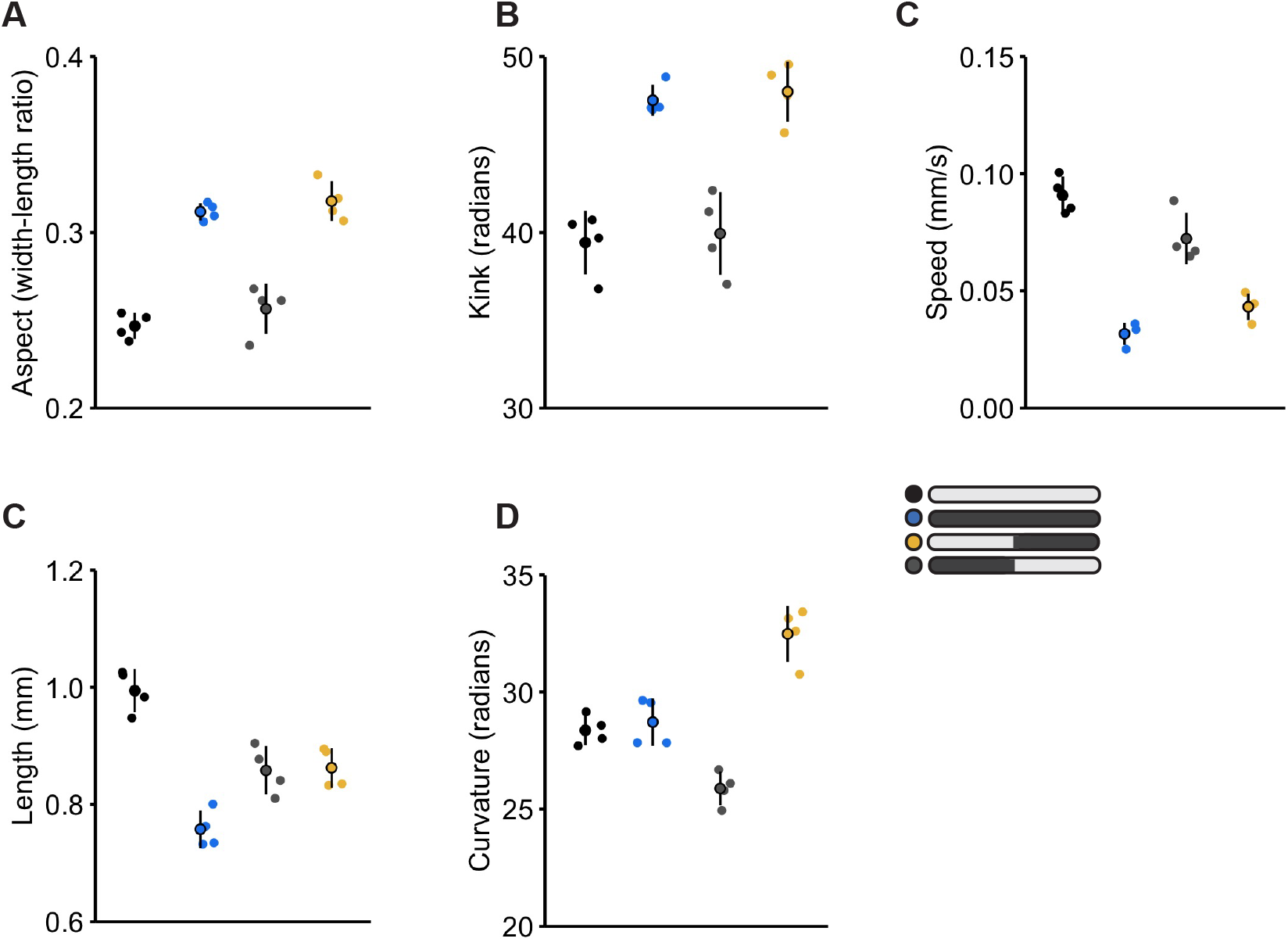
Pan-neuronally degrading and re-expressing dynein reveals distinct temporal functional windows for morphological and baseline locomotion features. The no-Auxin control group is depicted in black and continuous degradation group is depicted in blue for all panels. (A-C) Dynein is continuously required in neurons for normal kinked body posture, aspect-ratio, and movement direction bias. Degrading DHC-1 in neurons beginning at L4 (48 hrs post-hatch) induced impairment levels across these phenotypes similar to the continuous degradation control. Pan neuronal re-expression of DHC-1 at L4 (grey) rescued impairments in all three phenotypes. D) Re-expression of DHC-1 at L4 rescued speed before the onset mechanosensory stimuli, however only partially rescued speed deficits in the 5 min rest period. Degrading dynein at L4 (yellow) caused animals to exhibit lower speed throughout the behavioural paradigm that was similar to the continuous degradation control. E) Dynein is continuously required in neurons for animal length, but re-expression at L4 can partially rescue impairments F) Novel impairments in animal curvature occurred when dynein was re-expressed at L4. Beginning protein degradation at L4 caused animals to have a higher body curvature than the no-Auxin control, whereas re-expressing pan neuronal DHC-1 at L4 caused animals to exhibit a lower body curvature than controls.

We next investigated the phenotypic consequence of degrading and re-expressing dynein in neurons at an earlier time point in development. Re-expressing dynein during early post-embryonic development (starting Auxin exposure at L2) fully rescued impairments in response probability while starting degradation at L2 produced a similar level of impairment in response probability as the continuous degradation condition (**Fig. 6F**). Importantly, the lack of the novel hyperresponsive reversal probability phenotype that occurred when dynein was re-expressed at L4 suggests that for certain genes there will be crucial windows in development when re-expression must occur to avoid inducing alternative impairments. For reversal duration and speed, impairments in both phenotypes were fully rescued with re-expression starting at L2 (**Fig. 6G-H**). Earlier degradation did not induce impairments in reversal duration, suggesting the crucial functional window of DHC-1 for reversal duration is occurring prior to L2 but is not required throughout the lifespan (**Fig. 6G**). Earlier degradation did lower reversal speed (**Fig. 6H**), providing more evidence that DHC-1 is continuously required for this phenotype. Together, these results reveal many new roles for dynein in the developing and adult nervous system and illustrate the diversity of temporal functional windows that can be observed for a single gene (**Fig. 6E**).

### Comparison of temporal profiles reveals shared phenotypic disruptions and prioritizing principles of phenotypic reversibility

All neurodevelopmental disorder risk genes assessed here showed post-embryonic reversibility for at least one impaired phenotype, with several showing phenotypic reversibility relatively late in development (e.g. after stopping Auxin exposure at L4; **Fig. 8A**). Looking across all genes assessed, at least one phenotype within each phenotypic class (morphology, locomotion, mechanosensory responding, and learning) could be reversed later in life, suggesting a degree of flexibility in when multiple cellular processes can occur during development (**Fig. 8A**). However, not all phenotypes caused by the inactivation of a single gene could be reversed, even if reversibility was possible for other affected phenotypes (**Fig. 8A**). These results suggest that multiple phenotypic disruptions stemming from a single affected gene can show distinct windows of reversibility (**Fig. 8A**). In addition, there were also cases where the same organism-level phenotypic disruption (e.g. impaired mechanosensory responding) could be rescued later in development by re-expression of one of the genes, but not others (**Fig. 8A-C**). Based on our results, *EBF3●unc-3* is a promising candidate for further research as many impaired phenotypes could be reversed even when gene function was restored later stages of development (**Fig. 8A**), and re-expression did not induce novel phenotypic disruptions (**Fig. 2 & 3**). Further, identifying which phenotypes are most commonly affected across risk genes may provide insight into points of convergence in the mechanisms that are altered in neurodevelopmental disorders (**Fig. 8B**). We found that reversal probability metrics appeared to be both the most affected and reversible phenotypes across all genes assessed (**Fig. 8B & C**). This finding supports previous work from multiple model organisms which found that inactivating neurodevelopmental disorder risk genes commonly impaired habituation of response probability (Fenckova et al., 2019; McDiarmid et al., 2020) and that impairments in habituation caused by inactivation of the ASD risk gene ortholog neuroligin (*NLGN1/2/3/X•nlg-1*) could be partially rescued with adult re-expression (McDiarmid et al., 2020).

**Figure 8.**
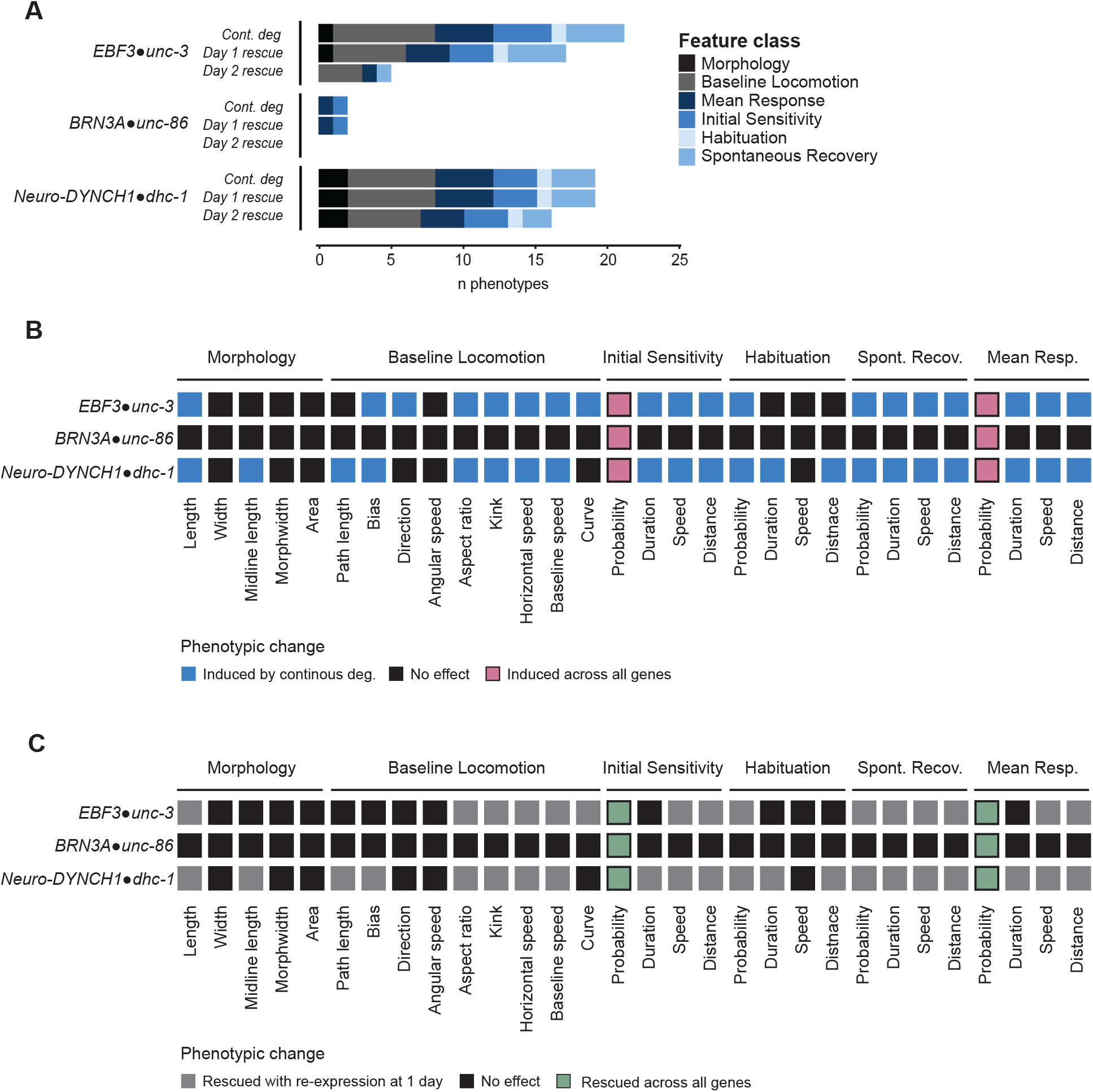
Comparison of temporal profiles reveals shared phenotypic disruptions and prioritizing principles of phenotypic reversibility. A) Number and kind of phenotypes that could be induced by continuously degrading each gene or reversed by re-expressing the gene 24h (L2 stage) or 48h (L4 stage) after synchronization (i.e. early or late in post-embryonic development). B) Heatmap showing the phenotypes affected by continuous degradation of each gene. Phenotypes observed in all 3 genes are highlighted in red. C) Heatmap showing the phenotypes that could be rescued with protein re-expression starting at L2 (24 hours post synchronization). Phenotypes that were reversible across all 3 genes are highlighted in green.

## Discussion

We systematically investigated the effect of degrading and re-expressing multiple neurodevelopmental disorder risk gene orthologs across a suite of morphological, locomotor, sensory, and learning phenotypes in thousands of freely behaving animals using our high-throughput machine vision tracking system. Taking advantage of the CRISPR-Cas9 AID system allowed us to test whether restoring protein levels through re-expression from the endogenous locus was sufficient for phenotypic rescue at multiple time points throughout development. We found that each gene displayed unique temporal functional windows and phenotypic profiles (**Fig. 8**). *DYNCH1●dhc-1* function is continuously essential for development and plays a specific role in adult mechanosensory behavior. The transcription factor *BRN3A●unc-86* can only reverse phenotypic disruptions early in post-embryonic development and is not required for adult function, suggesting a primarily developmental role in mechanosensory responding. The transcription factor *EBF3●unc-3* displayed a range of temporal requirements throughout the lifespan and can reverse multiple phenotypic disruptions later in life. In addition to the 3 genes tested in this study, previous findings from our lab provide the temporal requirements for another neurodevelopmental disorder risk gene, the synaptic cell adhesion molecule *NLGN1/2/3/4/X●nlg-1.* Adulthood re-expression of *NLGN1/2/3/4/X●nlg-1* can partially reverse impairments in habituation of response probability, however, once neuroligin has functioned to build a circuit capable of normal sensory processing it is no longer required in adulthood for normal short-term habituation (McDiarmid et al., 2020). Together, these results reveal a remarkable diversity in temporal phenotypic profiles across neurodevelopmental disorder risk genes that would be missed by approaches that focus on a single phenotype or developmental timepoint. The approach established in this study can be used to systematically assess the temporal requirements and phenotypic reversibility of neurodevelopmental disorder risk genes at an unprecedented throughput to prioritize risk genes for further assessment.

Using neuron-specific reversible protein degradation, we provide the first description of the role of dynein in behaviour across development. We found that continuous degradation of dynein in only neurons affected the majority (22/30) of the phenotypes assessed, including multiple morphology phenotypes. Interestingly, we found that both degrading or re-expressing dynein in neurons during early adulthood impaired habituation of mechanosensory response probability. Previous work from our lab has found habituation of response probability is affected by developmental stage such that habituation becomes deeper with age due to circuit rewiring and reduced sensitivity (Beck and Rankin, 1993; Rai and Rankin, 2007; Timbers et al., 2013). The impaired habituation phenotypes seen with both development and adult-specific dynein degradation could both stem from an immature nervous system, such that impairing protein function during early development temporarily impedes developmental processes from occurring whereas early adult inactivation freezes the nervous system in an immature state. Determining whether this neurodevelopmental freeze mechanism, or alternative, more complex mechanisms (e.g. changes in synaptic physiology) mediate these habituation impairments requires further study. Taken together these results reveal that dynein has several ongoing functions in the nervous system to modulate sensory and learning behaviors after it’s essential period in development.

Using high-throughput model systems to rapidly assess the temporal function and phenotypic reversibility of neurodevelopmental risk genes provides critical insight into the emerging principles that should be considered in future re-expression studies. Importantly, we found restoring protein expression in adulthood may induce novel phenotypes that were not observed when proteins were continuously degraded (e.g. altered habituation with neuron-specific re-expression of dynein). This finding reveals the importance of assessing a large number of morphological and behavioral phenotypes to ensure novel adverse phenotypes do not arise when gene function is restored later in development. These studies also offer a reminder to the diversity of functions of a single gene across development. Although a gene may play an important role in a phenotype under study, it also may have other functions that are not immediately obvious. Only by studying multiple phenotypes over the span of development can we begin to understand the breadth of what a given gene contributes to the organism. Future re-expression studies should not only aim to assess whether restoring gene function rescues the well-documented cell functions of the gene of interest, but also capture multiple organism-level phenotypes such as forms of learning or other behaviours commonly altered in neurodevelopmental disorders. Overall, our results suggest that earlier protein re-expression will almost always be better, yet identification of genes with longer reversibility windows and multiple reversible phenotypes in high-throughput model systems should be a key principle in prioritizing candidates for further study.

In addition, we found that time windows for when a gene is required and when re-expression can reverse impairments did not always align. While *EBF3●unc-3* showed reciprocal functional and reversibility windows for many affected phenotypes, we found other genes (e.g. *nlg-1* and *dhc-1*) where certain phenotypic impairments were not induced if the protein was degraded in early adulthood, but phenotypic rescue was possible if protein levels were restored at that same time point in development. In addition, while *EBF3●unc-3* and *BRN3A●unc-86* have relatively similar functions and previously described temporal functional windows, we identified stark differences in their reversibility profiles. Together, these findings highlight the need for systematic assessment of the reversibility windows of neurodevelopmental disorder risk genes across different developmental timepoints, even if there is prior indication of when the gene normally functions.

As the number of genes assessed increases, we may uncover patterns in the molecular attributes that enable certain genes to rescue impairments more broadly than others. For example, *EBF3●unc-3* is a putative pioneer transcription factor that may be able to more efficiently remodel chromatin and rewire transcriptional networks outside of its typical developmental window compared to other transcription factors. The approach developed in this study can be adapted to determine how gene reactivation reverses more complex behaviours in *C. elegans* and more conserved model systems. Information gained from high-throughput model organisms is increasingly valuable as they enable rapid assessment of the growing list of risk genes to gain insight into principles governing neurodevelopment and how a nervous system adapts to the re- introduction of a previously inactive protein.

## METHODS

### Animal maintenance

Prior to Auxin experiments, all strains were maintained on Petri plates containing Nematode Growth Medium (NGM) that were seeded with *Escherichia coli* (*E. coli*) strain OP50 following standard experimental procedures (Brenner, 1974). 96hr post-hatch hermaphrodite animals were used for all experiments.

### Auxin inducible degradation strain selection and ortholog identification

All human orthologs of all Auxin-inducible degradation strains available at the *CGC* were identified using the Alliance of Genome Resources ortholog prediction tool and OrthoList 2 (Agapite et al., 2020; Kim et al., 2018; McDiarmid et al., 2020). Auxin-inducible degradation strains for which the human ortholog corresponded to a known neurodevelopmental disorder risk gene (based on lists generated by recent large-scale sequencing studies and manual literature search (Belmadani et al., 2019; De Rubeis et al., 2014; McDiarmid et al., 2020; Satterstrom et al., 2020)) were selected for analysis. Note that throughout the manuscript the “*●*” symbol is used to denote the relationship between the human gene and *C. elegans* ortholog under study (e.g. *DYNCH1●dhc-1)*.

### Auxin plate preparation

Auxin administration was performed by transferring animals to bacteria-seeded NGM plates containing Auxin (McDiarmid et al., 2020; Zhang et al., 2015). To prepare Auxin plates, a 400 mM stock solution of Auxin indole-3- acetic acid (IAA) (Thermo Fisher, Alfa Aesar™ #A1055614) was created by dissolving Auxin in ethanol. Molten NGM was prepared and allowed to cool to approximately 50°C. The Auxin stock was then diluted into separate flasks of molten NGM agar to final concentrations of 0.025mM, 1mM, and 4mM. The NGM agar + Auxin mixture was then poured into Petri plates and allowed to dry in the dark for 72 hrs. Auxin plates were then seeded with 50 µl of *E. coli* OP50 liquid culture 48 hrs before use. All plates were stored in the dark at room temperature (20°C) in a temperature and humidity-controlled room (McDiarmid et al., 2020; Zhang et al., 2015).

### Population age-synchronization and Auxin administration

Age synchronization by egg lay was used to create the experimental groups for phenotypic analysis as previously described (McDiarmid et al., 2020; McDiarmid et al., 2020). For age synchronization, five gravid adults were placed on to either NGM or Auxin plates and allowed to lay eggs for 4 hours before removal (resulting in 50-100 animals per plate). For the development specific-degradation conditions, approximately 240 progeny were manually transferred from auxin plates onto 6 regular NGM plates (∼40 animals per plate) either 24 hrs (at L2), 48 hrs (at L4), or 72 hrs (at early adulthood) after synchronization (egg lay). For adult-specific degradation conditions, approximately 240 progeny were manually transferred from regular NGM plates onto 6 Auxin plates (∼40 animals per plate) 24 hrs (at L2), 48 hrs (at L4), or 72 hrs (at early adulthood) after synchronization. All plates remained in the dark other than when animals were being manually transferred to preserve the integrity of Auxin. 4-6 plates were run for each experimental condition.

### Behavioral paradigm and Multi-Worm Tracker phenotypic analysis

The Multi-Worm Tracker (MWT) was used for all behavioural tracking experiments (Swierczek et al., 2011). Each plate was subjected to the same short-term habituation behavioural paradigm (see results and **Fig. 1D**) that began with a 5 min period to allow worms to acclimate to being placed on the MWT. After acclimation, we collected data for an additional 5 min period to assess baseline locomotion and morphology features (**Fig. 1D**). Following this baseline period, thirty mechanosensory stimuli were administered to the side of the Petri plate using an automated push-solenoid at a 10 second inter-stimulus interval (**Fig. 1D**). These non-localized mechanosensory stimuli cause animals to perform a reversal response, where animals briefly crawl backwards before resuming forward locomotion (Rankin et al., 1990). We quantified multiple mechanosensory sensitivity and habituation learning phenotypes from these reversals which we have previously shown are mediated by genetically dissociable underlying mechanisms. After the 30^th^ stimulus, a 5 min rest period occurred which was followed by the administration of a final stimulus to assess short-term memory retention of habituation (spontaneous recovery, **Fig. 1D**). See The Multi-Worm Tracker user guide (https://sourceforge.net/projects/mwt/) or McDiarmid *et al*. 2020 (McDiarmid et al., 2020) for full description of all phenotypes. All testing occurred in a temperature and humidity-controlled room at approximately 20°C.

We used the MWT software (version 1.2.0.2) to delivery stimuli and acquire images (Swierczek et al., 2011), and Choreography software (version 1.3.0_r103552) to quantify phenotypes. Choreography filters “–shadowless”, “–minimum-move-body 2”, and “–minimum-time 20” were used to restrict analysis to animals that moved more than 2 body lengths and were tracked for 20 secs or longer. The “MeasureReversal” plug-in was used to identify animals that reversed within 1 sec of the mechanosensory stimulus being administered (Swierczek et al., 2011). Choreography output files were organized using custom R scripts which are freely available at [Github link will be inserted here]. All phenotypic features were pooled across the 4-6 plate replicates (each plate replicate capturing 40-100 animals) per strain. The mean of each condition was then compared using an unpaired t-test and Benjamini-Hochberg control of false discovery rate at 0.01. Figures were generated using the ggplot2 package in R (Wickham, 2016).

### Strains used

The following strains are available through the *Caenorhabditis Genetics Center* (*CGC*):

CA1200 *ieSi57[eft-3p::TIR1::mRuby::unc-54 3’UTR* + *cbr-unc-119(+)] II*
OH13988 *ieSi57[eft-3p::TIR1::mRuby::unc-54 3’UTR* + *cbr-unc-119(+)] II*; *unc-3(ot837[unc-3::mNeonGreen::AID]) X*
OH15227 *unc-86(ot893[unc-86::3xFlag::mNeonGreen::AID]) III*
CA1207 *dhc-1(ie28[dhc-1::degron::GFP]) I*
CA1210 *ieSi57[eft-3p::TIR1::mRuby::unc-54 3’UTR* + *cbr-unc-119(+)] II*; *dhc-1(ie28[dhc- 1::degron::GFP]) I*

The following strains were generated using standard genetic crosses:

VG937 *ieSi57[eft-3p::TIR1::mRuby::unc-54 3’UTR* + *cbr-unc-119(+)] II*; *unc-86(ot893[unc- 86::3xFlag::mNeonGreen::AID]) III*
VG946 *mizSi6[rab-3p::TIR1::unc-54 3’UTR* + *LoxP pmyo-2::GFP::unc-54 3’UTR prps27::NeoR::unc-54 3’UTR LoxP] V*; *dhc-1(ie28[dhc-1::degron::GFP]) I*

### Genotype confirmation

Successful crosses were determined through visual confirmation of fluorescent reporters as well as PCR-based genotyping using the following primers:

*TIR1* sequence

Forward: GACCGTAACTCCGTCTCC
Reverse: CGTTGGTGGTGATGATTTGAC

AID degron sequence

Forward: CCTAAAGATCCAGCCAAACC
Reverse: CTTCACGAACGCCGC

or

Forward: GATCCAGCCAAACCTCCGGC
Reverse: CTTCACGAACGCCGCCGC

## ACKNOWLEDGEMENTS

The authors thank Dr. Kota Mizumoto for the *rab-3p*::*TIR1* strain that was used for neuron-specific degradation and the *CGC* (funded by National Institute of Health Office of Research Infrastructure Programs, P40 OD010440) for other strains used within this research.

## AUTHOR CONTRIBUTIONS

Conceptualization: T.A.M and L.D.K. Methodology: T.A.M. Software and formal analysis: T.A.M. Investigation: L.D.K and T.A.M. Data curation: T.A.M. Writing-original draft preparation: L.D.K. and T.A.M. Writing- review and editing L.D.K, T.A.M, and C.H.R. Visualization: T.A.M. and L.D.K. Supervision: C.H.R. Funding acquisition: C.H.R.

## REFERENCES

Abrahams, B.S., Arking, D.E., Campbell, D.B., Mefford, H.C., Morrow, E.M., Weiss, L.A., Menashe, I., Wadkins, T., Banerjee-Basu, S., Packer, A., 2013. SFARI Gene 2.0: a community-driven knowledgebase for the autism spectrum disorders (ASDs). Mol. Autism 4, 36. https://doi.org/10.1186/2040-2392-4-36

Agapite, J., Albou, L.-P., Aleksander, S., Argasinska, J., Arnaboldi, V., Attrill, H., Bello, S.M., Blake, J.A., Blodgett, O., Bradford, Y.M., Bult, C.J., Cain, S., Calvi, B.R., Carbon, S., Chan, J., Chen, W.J., Cherry, J.M., Cho, J., Christie, K.R., Crosby, M.A., Pons, J. De, Dolan, M.E., Santos, G. dos, Dunn, B., Dunn, N., Eagle, A., Ebert, D., Engel, S.R., Fashena, D., Frazer, K., Gao, S., Gondwe, F., Goodman, J., Gramates, L.S., Grove, C.A., Harris, T., Harrison, M.-C., Howe, D.G., Howe, K.L., Jha, S., Kadin, J.A., Kaufman, T.C., Kalita, P., Karra, K., Kishore, R., Laulederkind, S., Lee, R., MacPherson, K.A., Marygold, S.J., Matthews, B., Millburn, G., Miyasato, S., Moxon, S., Mueller, H.-M., Mungall, C., Muruganujan, A., Mushayahama, T., Nash, R.S., Ng, P., Paulini, M., Perrimon, N., Pich, C., Raciti, D., Richardson, J.E., Russell, M., Gelbart, S.R., Ruzicka, L., Schaper, K., Shimoyama, M., Simison, M., Smith, C., Shaw, D.R., Shrivatsav, A., Skrzypek, M., Smith, J.R., Sternberg, P.W., Tabone, C.J., Thomas, P.D., Thota, J., Toro, S., Tomczuk, M., Tutaj, Marek, Tutaj, Monika, Urbano, J.-M., Auken, K. Van, Slyke, C.E. Van, Wang, S.-J., Weng, S., Westerfield, M., Williams, G., Wong, E.D., Wright, A., Yook, K., 2020. Alliance of Genome Resources Portal: unified model organism research platform. Nucleic Acids Res. 48, D650–D658. https://doi.org/10.1093/nar/gkz813

American Psychiatric Association, 2013. Diagnostic and Statistical Manual of Mental Disorders.

American Psychiatric Association. https://doi.org/10.1176/appi.books.9780890425596

Ardiel, E.L., McDiarmid, T.A., Timbers, T.A., Lee, K.C.Y., Safaei, J., Pelech, S.L., Rankin, C.H., 2018. Insights into the roles of CMK-1 and OGT-1 in interstimulus interval-dependent habituation in Caenorhabditis elegans. Proc. R. Soc. B Biol. Sci. 285. https://doi.org/10.1098/rspb.2018.2084

Ashley, G.E., Duong, T., Levenson, M.T., Martinez, M.A.Q., Johnson, L.C., Hibshman, J.D., Saeger, H.N., Palmisano, N.J., Doonan, R., Martinez-Mendez, R., Davidson, B.R., Zhang, W., Ragle, J.M., Medwig-Kinney, T.N., Sirota, S.S., Goldstein, B., Matus, D.Q., Dickinson, D.J., Reiner, D.J., Ward, J.D., 2021. An expanded auxin-inducible degron toolkit for Caenorhabditis elegans. Genetics 217. https://doi.org/10.1093/genetics/iyab006

Au, V., Li-Leger, E., Raymant, G., Flibotte, S., Chen, G., Martin, K., Fernando, L., Doell, C., Rosell, F.I., Wang, S., Edgley, M.L., Rougvie, A.E., Hutter, H., Moerman, D.G., 2019. CRISPR/Cas9 Methodology for the Generation of Knockout Deletions in Caenorhabditis elegans. G3&#58; Genes|Genomes|Genetics 9, 135–144. https://doi.org/10.1534/g3.118.200778

Badea, T.C., Cahill, H., Ecker, J., Hattar, S., Nathans, J., 2009. Distinct Roles of Transcription Factors Brn3a and Brn3b in Controlling the Development, Morphology, and Function of Retinal Ganglion Cells. Neuron 61, 852–864. https://doi.org/10.1016/j.neuron.2009.01.020

Beck, C.D.O., Rankin, C.H., 1993. Effects of aging on habituation in the nematode Caenorhabditis elegans. Behav. Processes 28, 145–163. https://doi.org/10.1016/0376-6357(93)90088-9

Belmadani, M., Jacobson, M., Holmes, N., Phan, M., Nguyen, T., Pavlidis, P., Rogic, S., 2019. VariCarta: A Comprehensive Database of Harmonized Genomic Variants Found in Autism Spectrum Disorder Sequencing Studies. Autism Res. 12, 1728–1736. https://doi.org/10.1002/aur.2236

Boyle, C.A., Boulet, S., Schieve, L.A., Cohen, R.A., Blumberg, S.J., Yeargin-Allsopp, M., Visser, S., Kogan, M.D., 2011. Trends in the Prevalence of Developmental Disabilities in US Children, 1997-2008. Pediatrics 127, 1034–1042. https://doi.org/10.1542/peds.2010-2989

Brenner, S., 1974. The genetics of Caenorhabditis elegans. Genetics 77, 71–94.

Chao, H.-T., Davids, M., Burke, E., Pappas, J.G., Rosenfeld, J.A., McCarty, A.J., Davis, T., Wolfe, L., Toro, C., Tifft, C., Xia, F., Stong, N., Johnson, T.K., Warr, C.G., Yamamoto, S., Adams, D.R., Markello, T.C., Gahl, W.A., Bellen, H.J., Wangler, M.F., Malicdan, M.C. V., Adams, D.R., Adams, C.J., Alejandro, M.E., Allard, P., Ashley, E.A., Bacino, C.A., Balasubramanyam, A., Barseghyan, H., Beggs, A.H., Bellen, H.J., Bernstein, J.A., Bick, D.P., Birch, C.L., Boone, B.E., Briere, L.C., Brown, D.M., Brush, M., Burrage, L.C., Chao, K.R., Clark, G.D., Cogan, J.D., Cooper, C.M., Craigen, W.J., Davids, M., Dayal, J.G., Dell’Angelica, E.C., Dhar, S.U., Dipple, K.M., Donnell-Fink, L.A., Dorrani, N., Dorset, D.C., Draper, D.D., Dries, A.M., Eckstein, D.J., Emrick, L.T., Eng, C.M., Esteves, C., Estwick, T., Fisher, P.G., Frisby, T.S., Frost, K., Gahl, W.A., Gartner, V., Godfrey, R.A., Goheen, M., Golas, G.A., Goldstein, D.B., Gordon, M. “Gracie” G., Gould, S.E., Gourdine, J.-P.F., Graham, B.H., Groden, C.A., Gropman, A.L., Hackbarth, M.E., Haendel, M., Hamid, R., Hanchard, N.A., Handley, L.H., Hardee, I., Herzog, M.R., Holm, I.A., Howerton, E.M., Jacob, H.J., Jain, M., Jiang, Y., Johnston, J.M., Jones, A.L., Koehler, A.E., Koeller, D.M., Kohane, I.S., Kohler, J.N., Krasnewich, D.M., Krieg, E.L., Krier, J.B., Kyle, J.E., Lalani, S.R., Latham, L., Latour, Y.L., Lau, C.C., Lazar, J., Lee, B.H., Lee, H., Lee, P.R., Levy, S.E., Levy, D.J., Lewis, R.A., Liebendorder, A.P., Lincoln, S.A., Loomis, C.R., Loscalzo, J., Maas, R.L., Macnamara, E.F., MacRae, C.A., Maduro, V. V., Malicdan, M.C. V., Mamounas, L.A., Manolio, T.A., Markello, T.C., Mashid, A.S., Mazur, P., McCarty, A.J., McConkie-Rosell, A., McCray, A.T., Metz, T.O., Might, M., Moretti, P.M., Mulvihill, J.J., Murphy, J.L., Muzny, D.M., Nehrebecky, M.E., Nelson, S.F., Newberry, J.S., Newman, J.H., Nicholas, S.K., Novacic, D., Orange, J.S., Pallais, J.C., Palmer, C.G.S., Papp, J.C., Pena, L.D.M., Phillips, J.A., Posey, J.E., Postlethwait, J.H., Potocki, L., Pusey, B.N., Ramoni, R.B., Rodan, L.H., Sadozai, S., Schaffer, K.E., Schoch, K., Schroeder, M.C., Scott, D.A., Sharma, P., Shashi, V., Silverman, E.K., Sinsheimer, J.S., Soldatos, A.G., Spillmann, R.C., Splinter, K., Stoler, J.M., Stong, N., Strong, K.A., Sullivan, J.A., Sweetser, D.A., Thomas, S.P., Tift, C.J., Tolman, N.J., Toro, C., Tran, A.A., Valivullah, Z.M., Vilain, E., Waggott, D.M., Wahl, C.E., Walley, N.M., Walsh, C.A., Wangler, M.F., Warburton, M., Ward, P.A., Waters, K.M., Webb-Robertson, B.-J.M., Weech, A.A., Westerfield, M., Wheeler, M.T., Wise, A.L., Worthe, L.A., Worthey, E.A., Yamamoto, S., Yang, Y., Yu, G., Zornio, P.A., 2017. A Syndromic Neurodevelopmental Disorder Caused by De Novo Variants in EBF3. Am. J. Hum. Genet. 100, 128–137. https://doi.org/10.1016/j.ajhg.2016.11.018

Cianfrocco, M.A., DeSantis, M.E., Leschziner, A.E., Reck-Peterson, S.L., 2015. Mechanism and Regulation of Cytoplasmic Dynein. Annu. Rev. Cell Dev. Biol. 31, 83–108. https://doi.org/10.1146/annurev-cellbio-100814-125438

Creson, T.K., Rojas, C., Hwaun, E., Vaissiere, T., Kilinc, M., Jimenez-Gomez, A., Holder, J.L., Tang, J., Colgin, L.L., Miller, C.A., Rumbaugh, G., 2019. Re-expression of SynGAP protein in adulthood improves translatable measures of brain function and behavior. Elife 8. https://doi.org/10.7554/eLife.46752

de la Torre-Ubieta, L., Won, H., Stein, J.L., Geschwind, D.H., 2016. Advancing the understanding of autism disease mechanisms through genetics. Nat. Med. 22, 345–361. https://doi.org/10.1038/nm.4071

De Rubeis, S., He, X., Goldberg, A.P., Poultney, C.S., Samocha, K., Ercument Cicek, A., Kou, Y., Liu, L., Fromer, M., Walker, S., Singh, T., Klei, L., Kosmicki, J., Fu, S.-C., Aleksic, B., Biscaldi, M., Bolton, P.F., Brownfeld, J.M., Cai, J., Campbell, N.G., Carracedo, A., Chahrour, M.H., Chiocchetti, A.G., Coon, H., Crawford, E.L., Crooks, L., Curran, S.R., Dawson, G., Duketis, E., Fernandez, B.A., Gallagher, L., Geller, E., Guter, S.J., Sean Hill, R., Ionita-Laza, I., Jimenez Gonzalez, P., Kilpinen, H., Klauck, S.M., Kolevzon, A., Lee, I., Lei, J., Lehtimäki, T., Lin, C.-F., Ma’ayan, A., Marshall, C.R., McInnes, A.L., Neale, B., Owen, M.J., Ozaki, N., Parellada, M., Parr, J.R., Purcell, S., Puura, K., Rajagopalan, D., Rehnström, K., Reichenberg, A., Sabo, A., Sachse, M., Sanders, S.J., Schafer, C., Schulte-Rüther, M., Skuse, D., Stevens, C., Szatmari, P., Tammimies, K., Valladares, O., Voran, A., Wang, L.-S., Weiss, L.A., Jeremy Willsey, A., Yu, T.W., Yuen, R.K.C., Cook, E.H., Freitag, C.M., Gill, M., Hultman, C.M., Lehner, T., Palotie, A., Schellenberg, G.D., Sklar, P., State, M.W., Sutcliffe, J.S., Walsh, C.A., Scherer, S.W., Zwick, M.E., Barrett, J.C., Cutler, D.J., Roeder, K., Devlin, B., Daly, M.J., Buxbaum, J.D., 2014. Synaptic, transcriptional and chromatin genes disrupted in autism. Nature 515, 209–215. https://doi.org/10.1038/nature13772

Deciphering Developmental Disorders Study, 2015. Large-scale discovery of novel genetic causes of developmental disorders. Nature 519, 223–228. https://doi.org/10.1038/nature14135

Dickinson, D.J., Goldstein, B., 2016. CRISPR-Based Methods for Caenorhabditis elegans Genome Engineering. Genetics 202, 885–901. https://doi.org/10.1534/genetics.115.182162

Ehninger, D., Li, W., Fox, K., Stryker, M.P., Silva, A.J., 2008. Reversing Neurodevelopmental Disorders in Adults. Neuron 60, 950–960. https://doi.org/10.1016/j.neuron.2008.12.007

Fenckova, M., Blok, L.E.R., Asztalos, L., Goodman, D.P., Cizek, P., Singgih, E.L., Glennon, J.C., IntHout, J., Zweier, C., Eichler, E.E., von Reyn, C.R., Bernier, R.A., Asztalos, Z., Schenck, A., 2019. Habituation Learning is a Widely Affected Mechanism in Drosophila Models of Intellectual Disability and Autism Spectrum Disorders. Biol. Psychiatry. https://doi.org/10.1016/j.biopsych.2019.04.029

Feng, W., Li, Y., Dao, P., Aburas, J., Islam, P., Elbaz, B., Kolarzyk, A., Brown, A.E., Kratsios, P., 2020. A terminal selector prevents a Hox transcriptional switch to safeguard motor neuron identity throughout life. Elife 9. https://doi.org/10.7554/eLife.50065

Gao, Y., Irvine, E.E., Eleftheriadou, I., Naranjo, C.J., Hearn-Yeates, F., Bosch, L., Glegola, J.A., Murdoch, L., Czerniak, A., Meloni, I., Renieri, A., Kinali, M., Mazarakis, N.D., 2020. Gene replacement ameliorates deficits in mouse and human models of cyclin-dependent kinase- like 5 disorder. Brain 143, 811–832. https://doi.org/10.1093/brain/awaa028

Green, S.A., Hernandez, L., Lawrence, K.E., Liu, J., Tsang, T., Yeargin, J., Cummings, K., Laugeson, E., Dapretto, M., Bookheimer, S.Y., 2019. Distinct Patterns of Neural Habituation and Generalization in Children and Adolescents With Autism With Low and High Sensory Overresponsivity. Am. J. Psychiatry 176, 1010–1020. https://doi.org/10.1176/appi.ajp.2019.18121333

Green, S.A., Hernandez, L., Tottenham, N., Krasileva, K., Bookheimer, S.Y., Dapretto, M., 2015. Neurobiology of Sensory Overresponsivity in Youth With Autism Spectrum Disorders. JAMA Psychiatry 72, 778. https://doi.org/10.1001/jamapsychiatry.2015.0737

Guy, J., Gan, J., Selfridge, J., Cobb, S., Bird, A., 2007. Reversal of Neurological Defects in a Mouse Model of Rett Syndrome. Science (80-.). 315, 1143–1147. https://doi.org/10.1126/science.1138389

Hamill, D.R., Severson, A.F., Carter, J.C., Bowerman, B., 2002. Centrosome Maturation and Mitotic Spindle Assembly in C. elegans Require SPD-5, a Protein with Multiple Coiled-Coil Domains. Dev. Cell 3, 673–684. https://doi.org/10.1016/S1534-5807(02)00327-1

Harada, A., Takei, Y., Kanai, Y., Tanaka, Y., Nonaka, S., Hirokawa, N., 1998. Golgi Vesiculation and Lysosome Dispersion in Cells Lacking Cytoplasmic Dynein. J. Cell Biol. 141, 51–59. https://doi.org/10.1083/jcb.141.1.51

Howell, A.M., Rose, A.M., 1990. Essential genes in the hDf6 region of chromosome I in Caenorhabditis elegans. Genetics 126, 583–92.

Huang, E.J., Liu, W., Fritzsch, B., Bianchi, L.M., Reichardt, L.F., Xiang, M., 2001. Brn3a is a transcriptional regulator of soma size, target field innervation and axon pathfinding of inner ear sensory neurons. Development 128, 2421–32.

Husson, Steven J., Steuer Costa, Wagner., Schmitt, Cornelia., Gottschalk, A., 2012. Keeping track of worm trackers (September 10, 2012), in: The C. elegans Research Community (Ed.), WormBook. https://doi.org/doi/10.1895/wormbook.1.150.1

Iakoucheva, L.M., Muotri, A.R., Sebat, J., 2019. Getting to the Cores of Autism. Cell 178, 1287– 1298. https://doi.org/10.1016/j.cell.2019.07.037

Iossifov, I., O’Roak, B.J., Sanders, S.J., Ronemus, M., Krumm, N., Levy, D., Stessman, H.A., Witherspoon, K.T., Vives, L., Patterson, K.E., Smith, J.D., Paeper, B., Nickerson, D.A., Dea, J., Dong, S., Gonzalez, L.E., Mandell, J.D., Mane, S.M., Murtha, M.T., Sullivan, C.A., Walker, M.F., Waqar, Z., Wei, L., Willsey, A.J., Yamrom, B., Lee, Y., Grabowska, E., Dalkic, E., Wang, Z., Marks, S., Andrews, P., Leotta, A., Kendall, J., Hakker, I., Rosenbaum, J., Ma, B., Rodgers, L., Troge, J., Narzisi, G., Yoon, S., Schatz, M.C., Ye, K., McCombie, W.R., Shendure, J., Eichler, E.E., State, M.W., Wigler, M., 2014. The contribution of de novo coding mutations to autism spectrum disorder. Nature 515, 216–221. https://doi.org/10.1038/nature13908

Jin, X., Simmons, S.K., Guo, A., Shetty, A.S., Ko, M., Nguyen, L., Jokhi, V., Robinson, E., Oyler, P., Curry, N., Deangeli, G., Lodato, S., Levin, J.Z., Regev, A., Zhang, F., Arlotta, P., 2020. In vivo Perturb-Seq reveals neuronal and glial abnormalities associated with autism risk genes. Science (80-.). 370, eaaz6063. https://doi.org/10.1126/science.aaz6063

Kaletta, T., Hengartner, M.O., 2006. Finding function in novel targets: C. elegans as a model organism. Nat. Rev. Drug Discov. 5, 387–399. https://doi.org/10.1038/nrd2031

Kavšek, M., 2004. Predicting later IQ from infant visual habituation and dishabituation: A meta- analysis. J. Appl. Dev. Psychol. 25, 369–393. https://doi.org/10.1016/j.appdev.2004.04.006

Kepler, L.D., McDiarmid, T.A., Rankin, C.H., 2020. Habituation in high-throughput genetic model organisms as a tool to investigate the mechanisms of neurodevelopmental disorders. Neurobiol. Learn. Mem. 171, 107208. https://doi.org/10.1016/j.nlm.2020.107208

Kim, W., Underwood, R.S., Greenwald, I., Shaye, D.D., 2018. OrthoList 2: A New Comparative Genomic Analysis of Human and Caenorhabditis elegans Genes. Genetics 210, 445–461. https://doi.org/10.1534/genetics.118.301307

Kindt, K.S., Quast, K.B., Giles, A.C., De, S., Hendrey, D., Nicastro, I., Rankin, C.H., Schafer, W.R., 2007. Dopamine Mediates Context-Dependent Modulation of Sensory Plasticity in C. elegans. Neuron 55, 662–676. https://doi.org/10.1016/j.neuron.2007.07.023

Kleinhans, N.M., Johnson, L.C., Richards, T., Mahurin, R., Greenson, J., Dawson, G., Aylward, E., 2009. Reduced Neural Habituation in the Amygdala and Social Impairments in Autism Spectrum Disorders. Am. J. Psychiatry 166, 467–475. https://doi.org/10.1176/appi.ajp.2008.07101681

Kratsios, P., Kerk, S.Y., Catela, C., Liang, J., Vidal, B., Bayer, E.A., Feng, W., De La Cruz, E.D., Croci, L., Consalez, G.G., Mizumoto, K., Hobert, O., 2017. An intersectional gene regulatory strategy defines subclass diversity of C. elegans motor neurons. Elife 6. https://doi.org/10.7554/eLife.25751

Levitan, D., Doyle, T.G., Brousseau, D., Lee, M.K., Thinakaran, G., Slunt, H.H., Sisodia, S.S., Greenwald, I., 1996. Assessment of normal and mutant human presenilin function in Caenorhabditis elegans. Proc. Natl. Acad. Sci. 93, 14940–14944. https://doi.org/10.1073/pnas.93.25.14940

Li, Y., Osuma, A., Correa, E., Okebalama, M.A., Dao, P., Gaylord, O., Aburas, J., Islam, P., Brown, A.E., Kratsios, P., 2020. Establishment and maintenance of motor neuron identity via temporal modularity in terminal selector function. Elife 9. https://doi.org/10.7554/eLife.59464

Lopes, F., Soares, G., Gonçalves-Rocha, M., Pinto-Basto, J., Maciel, P., 2017. Whole Gene Deletion of EBF3 Supporting Haploinsufficiency of This Gene as a Mechanism of Neurodevelopmental Disease. Front. Genet. 8. https://doi.org/10.3389/fgene.2017.00143

Mains, P.E., Sulston, I.A., Wood, W.B., 1990. Dominant maternal-effect mutations causing embryonic lethality in Caenorhabditis elegans. Genetics 125, 351–69.

Massa, J., O’Desky, I.H., 2012. Impaired Visual Habituation in Adults With ADHD. J. Atten. Disord. 16, 553–561. https://doi.org/10.1177/1087054711423621

McDiarmid, T. A., Au, V., Loewen, A.D., Liang, J., Mizumoto, K., Moerman, D.G., Rankin, C.H., 2018. CRISPR-Cas9 human gene replacement and phenomic characterization in Caenorhabditis elegans to understand the functional conservation of human genes and decipher variants of uncertain significance. Dis. Model. Mech. 11. https://doi.org/10.1242/dmm.036517

McDiarmid, T. A., Belmadani, M., Liang, J., Meili, F., Mathews, E.A., Mullen, G.P., Hendi, A., Wong, W.-R., Rand, J.B., Mizumoto, K., Haas, K., Pavlidis, P., Rankin, C.H., 2020. Systematic phenomics analysis of autism-associated genes reveals parallel networks underlying reversible impairments in habituation. Proc. Natl. Acad. Sci. 117, 656–667. https://doi.org/10.1073/pnas.1912049116

McDiarmid, T.A., Bernardos, A.C., Rankin, C.H., 2017. Habituation is altered in neuropsychiatric disorders—A comprehensive review with recommendations for experimental design and analysis. Neurosci. Biobehav. Rev. 80, 286–305. https://doi.org/10.1016/j.neubiorev.2017.05.028

McDiarmid, T.A, Kepler, L.D., Rankin, C.H., 2020. Auxin does not affect a suite of morphological or behavioral phenotypes in two wild-type C. elegans strains. microPublication Biol. 2020. https://doi.org/10.17912/micropub.biology.000307

McDiarmid, T.A., Yu, A.J., Rankin, C.H., 2019. Habituation Is More Than Learning to Ignore: Multiple Mechanisms Serve to Facilitate Shifts in Behavioral Strategy. BioEssays 41, 1900077. https://doi.org/10.1002/bies.201900077

McDiarmid, T. A., Yu, A.J., Rankin, C.H., 2018. Beyond the response-High throughput behavioral analyses to link genome to phenome in Caenorhabditis elegans. Genes, Brain Behav. 17, e12437. https://doi.org/10.1111/gbb.12437

Mei, Y., Monteiro, P., Zhou, Y., Kim, J.-A., Gao, X., Fu, Z., Feng, G., 2016. Adult restoration of Shank3 expression rescues selective autistic-like phenotypes. Nature 530, 481–484. https://doi.org/10.1038/nature16971

Nance, J., Frøkjær-Jensen, C., 2019. The Caenorhabditis elegans Transgenic Toolbox. Genetics 212, 959–990. https://doi.org/10.1534/genetics.119.301506

Nishimura, K., Fukagawa, T., Takisawa, H., Kakimoto, T., Kanemaki, M., 2009. An auxin-based degron system for the rapid depletion of proteins in nonplant cells. Nat. Methods 6, 917– 22. https://doi.org/10.1038/nmeth.1401

Parikshak, N.N., Luo, R., Zhang, A., Won, H., Lowe, J.K., Chandran, V., Horvath, S., Geschwind, D.H., 2013. Integrative Functional Genomic Analyses Implicate Specific Molecular Pathways and Circuits in Autism. Cell 155, 1008–1021. https://doi.org/10.1016/j.cell.2013.10.031

Post, K.L., Belmadani, M., Ganguly, P., Meili, F., Dingwall, R., McDiarmid, T.A., Meyers, W.M., Herrington, C., Young, B.P., Callaghan, D.B., Rogic, S., Edwards, M., Niciforovic, A., Cau, A., Rankin, C.H., O’Connor, T.P., Bamji, S.X., Loewen, C.J.R., Allan, D.W., Pavlidis, P., Haas, K., 2020. Multi-model functionalization of disease-associated PTEN missense mutations identifies multiple molecular mechanisms underlying protein dysfunction. Nat. Commun. 11, 2073. https://doi.org/10.1038/s41467-020-15943-0

Prasad, B., Karakuzu, O., Reed, R.R., Cameron, S., 2008. unc-3-dependent repression of specific motor neuron fates in Caenorhabditis elegans. Dev. Biol. 323, 207–215. https://doi.org/10.1016/j.ydbio.2008.08.029

Prasad, B.C., Ye, B., Zackhary, R., Schrader, K., Seydoux, G., Reed, R.R., 1998. unc-3, a gene required for axonal guidance in Caenorhabditis elegans, encodes a member of the O/E family of transcription factors. Development 125, 1561–8.

Rai, S., Rankin, C.H., 2007. Critical and sensitive periods for reversing the effects of mechanosensory deprivation on behavior, nervous system, and development inCaenorhabditis elegans. Dev. Neurobiol. 67, 1443–1456. https://doi.org/10.1002/dneu.20522

Randlett, O., Haesemeyer, M., Forkin, G., Shoenhard, H., Schier, A.F., Engert, F., Granato, M., 2019. Distributed Plasticity Drives Visual Habituation Learning in Larval Zebrafish. Curr. Biol. 29, 1337–1345.e4. https://doi.org/10.1016/j.cub.2019.02.039

Rankin, C.H., Abrams, T., Barry, R.J., Bhatnagar, S., Clayton, D.F., Colombo, J., Coppola, G., Geyer, M.A., Glanzman, D.L., Marsland, S., McSweeney, F.K., Wilson, D.A., Wu, C.-F., Thompson, R.F., 2009. Habituation revisited: An updated and revised description of the behavioral characteristics of habituation. Neurobiol. Learn. Mem. 92, 135–138. https://doi.org/10.1016/j.nlm.2008.09.012

Rankin, C.H., Beck, C.D.O., Chiba, C.M., 1990. Caenorhabditis elegans: A new model system for the study of learning and memory. Behav. Brain Res. 37, 89–92. https://doi.org/10.1016/0166-4328(90)90074-O

Robinson, J.T., Wojcik, E.J., Sanders, M.A., McGrail, M., Hays, T.S., 1999. Cytoplasmic dynein is required for the nuclear attachment and migration of centrosomes during mitosis in Drosophila. J. Cell Biol. 146, 597–608. https://doi.org/10.1083/jcb.146.3.597

Sanders, S.J., He, X., Willsey, A.J., Ercan-Sencicek, A.G., Samocha, K.E., Cicek, A.E., Murtha, M.T., Bal, V.H., Bishop, S.L., Dong, S., Goldberg, A.P., Jinlu, C., Keaney, J.F., Klei, L., Mandell, J.D., Moreno-De-Luca, D., Poultney, C.S., Robinson, E.B., Smith, L., Solli- Nowlan, T., Su, M.Y., Teran, N.A., Walker, M.F., Werling, D.M., Beaudet, A.L., Cantor, R.M., Fombonne, E., Geschwind, D.H., Grice, D.E., Lord, C., Lowe, J.K., Mane, S.M., Martin, D.M., Morrow, E.M., Talkowski, M.E., Sutcliffe, J.S., Walsh, C.A., Yu, T.W., Ledbetter, D.H., Martin, C.L., Cook, E.H., Buxbaum, J.D., Daly, M.J., Devlin, B., Roeder, K., State, M.W., 2015. Insights into Autism Spectrum Disorder Genomic Architecture and Biology from 71 Risk Loci. Neuron 87, 1215–1233. https://doi.org/10.1016/j.neuron.2015.09.016

Sanders, S.J., Sahin, M., Hostyk, J., Thurm, A., Jacquemont, S., Avillach, P., Douard, E., Martin, C.L., Modi, M.E., Moreno-De-Luca, A., Raznahan, A., Anticevic, A., Dolmetsch, R., Feng, G., Geschwind, D.H., Glahn, D.C., Goldstein, D.B., Ledbetter, D.H., Mulle, J.G., Pasca, S.P., Samaco, R., Sebat, J., Pariser, A., Lehner, T., Gur, R.E., Bearden, C.E., 2019. A framework for the investigation of rare genetic disorders in neuropsychiatry. Nat. Med. 25, 1477–1487. https://doi.org/10.1038/s41591-019-0581-5

Satterstrom, F.K., Kosmicki, J.A., Wang, J., Breen, M.S., De Rubeis, S., An, J.-Y., Peng, M., Collins, R., Grove, J., Klei, L., Stevens, C., Reichert, J., Mulhern, M.S., Artomov, M., Gerges, S., Sheppard, B., Xu, X., Bhaduri, A., Norman, U., Brand, H., Schwartz, G., Nguyen, R., Guerrero, E.E., Dias, C., Betancur, C., Cook, E.H., Gallagher, L., Gill, M., Sutcliffe, J.S., Thurm, A., Zwick, M.E., Børglum, A.D., State, M.W., Cicek, A.E., Talkowski, M.E., Cutler, D.J., Devlin, B., Sanders, S.J., Roeder, K., Daly, M.J., Buxbaum, J.D., Aleksic, B., Anney, R., Barbosa, M., Bishop, S., Brusco, A., Bybjerg-Grauholm, J., Carracedo, A., Chan, M.C.Y., Chiocchetti, A.G., Chung, B.H.Y., Coon, H., Cuccaro, M.L., Curró, A., Dalla Bernardina, B., Doan, R., Domenici, E., Dong, S., Fallerini, C., Fernández- Prieto, M., Ferrero, G.B., Freitag, C.M., Fromer, M., Gargus, J.J., Geschwind, D., Giorgio, E., González-Peñas, J., Guter, S., Halpern, D., Hansen-Kiss, E., He, X., Herman, G.E., Hertz-Picciotto, I., Hougaard, D.M., Hultman, C.M., Ionita-Laza, I., Jacob, S., Jamison, J., Jugessur, A., Kaartinen, M., Knudsen, G.P., Kolevzon, A., Kushima, I., Lee, S.L., Lehtimäki, T., Lim, E.T., Lintas, C., Lipkin, W.I., Lopergolo, D., Lopes, F., Ludena, Y., Maciel, P., Magnus, P., Mahjani, B., Maltman, N., Manoach, D.S., Meiri, G., Menashe, I., Miller, J., Minshew, N., Montenegro, E.M.S., Moreira, D., Morrow, E.M., Mors, O., Mortensen, P.B., Mosconi, M., Muglia, P., Neale, B.M., Nordentoft, M., Ozaki, N., Palotie, A., Parellada, M., Passos-Bueno, M.R., Pericak-Vance, M., Persico, A.M., Pessah, I., Puura, K., Reichenberg, A., Renieri, A., Riberi, E., Robinson, E.B., Samocha, K.E., Sandin, S., Santangelo, S.L., Schellenberg, G., Scherer, S.W., Schlitt, S., Schmidt, R., Schmitt, L., Silva, I.M.W., Singh, T., Siper, P.M., Smith, M., Soares, G., Stoltenberg, C., Suren, P., Susser, E., Sweeney, J., Szatmari, P., Tang, L., Tassone, F., Teufel, K., Trabetti, E., Trelles, M. del P., Walsh, C.A., Weiss, L.A., Werge, T., Werling, D.M., Wigdor, E.M., Wilkinson, E., Willsey, A.J., Yu, T.W., Yu, M.H.C., Yuen, R., Zachi, E., Agerbo, E., Als, T.D., Appadurai, V., Bækvad-Hansen, M., Belliveau, R., Buil, A., Carey, C.E., Cerrato, F., Chambert, K., Churchhouse, C., Dalsgaard, S., Demontis, D., Dumont, A., Goldstein, J., Hansen, C.S., Hauberg, M.E., Hollegaard, M. V., Howrigan, D.P., Huang, H., Maller, J., Martin, A.R., Martin, J., Mattheisen, M., Moran, J., Pallesen, J., Palmer, D.S., Pedersen, C.B., Pedersen, M.G., Poterba, T., Poulsen, J.B., Ripke, S., Schork, A.J., Thompson, W.K., Turley, P., Walters, R.K., 2020. Large-Scale Exome Sequencing Study Implicates Both Developmental and Functional Changes in the Neurobiology of Autism. Cell. https://doi.org/10.1016/j.cell.2019.12.036

Schmid, S., Wilson, D.A., Rankin, C.H., 2015. Habituation mechanisms and their importance for cognitive function. Front. Integr. Neurosci. 8, 97. https://doi.org/10.3389/fnint.2014.00097

Serrano-Saiz, E., Leyva-Díaz, E., De La Cruz, E., Hobert, O., 2018. BRN3-type POU Homeobox Genes Maintain the Identity of Mature Postmitotic Neurons in Nematodes and Mice. Curr. Biol. 28, 2813–2823.e2. https://doi.org/10.1016/j.cub.2018.06.045

Sleven, H., Welsh, S.J., Yu, J., Churchill, M.E.A., Wright, C.F., Henderson, A., Horvath, R., Rankin, J., Vogt, J., Magee, A., McConnell, V., Green, A., King, M.D., Cox, H., Armstrong, L., Lehman, A., Nelson, T.N., Williams, J., Clouston, P., Hagman, J., Németh, A.H., 2017. De Novo Mutations in EBF3 Cause a Neurodevelopmental Syndrome. Am. J. Hum. Genet. 100, 138–150. https://doi.org/10.1016/j.ajhg.2016.11.020

Swierczek, N.A., Giles, A.C., Rankin, C.H., Kerr, R.A., 2011. High-throughput behavioral analysis in C. elegans 8. https://doi.org/10.1038/nmeth.1625

Sze, J.Y., Ruvkun, G., 2003. Activity of the Caenorhabditis elegans UNC-86 POU transcription factor modulates olfactory sensitivity. Proc. Natl. Acad. Sci. 100, 9560–9565. https://doi.org/10.1073/pnas.1530752100

Tanaka, A.J., Cho, M.T., Willaert, R., Retterer, K., Zarate, Y.A., Bosanko, K., Stefans, V., Oishi, K., Williamson, A., Wilson, G.N., Basinger, A., Barbaro-Dieber, T., Ortega, L., Sorrentino, S., Gabriel, M.K., Anderson, I.J., Sacoto, M.J.G., Schnur, R.E., Chung, W.K., 2017. De novo variants in EBF3 are associated with hypotonia, developmental delay, intellectual disability, and autism. Mol. Case Stud. 3, a002097. https://doi.org/10.1101/mcs.a002097

Thompson, R.F., Spencer, W.A., 1966. Habituation: A model phenomenon for the study of neuronal substrates of behavior. Psychol. Rev. 73, 16–43. https://doi.org/10.1037/h0022681

Timbers, T.A., Giles, A.C., Ardiel, E.L., Kerr, R.A., Rankin, C.H., 2013. Intensity discrimination deficits cause habituation changes in middle-aged Caenorhabditis elegans. Neurobiol. Aging 34, 621–631. https://doi.org/10.1016/j.neurobiolaging.2012.03.016

Vissers, L.E.L.M., Gilissen, C., Veltman, J.A., 2016. Genetic studies in intellectual disability and related disorders. Nat. Rev. Genet. 17, 9–18. https://doi.org/10.1038/nrg3999

Vogel-Ciernia, A., Matheos, D.P., Barrett, R.M., Kramár, E.A., Azzawi, S., Chen, Y., Magnan, C.N., Zeller, M., Sylvain, A., Haettig, J., Jia, Y., Tran, A., Dang, R., Post, R.J., Chabrier, M., Babayan, A.H., Wu, J.I., Crabtree, G.R., Baldi, P., Baram, T.Z., Lynch, G., Wood, M.A., 2013. The neuron-specific chromatin regulatory subunit BAF53b is necessary for synaptic plasticity and memory. Nat. Neurosci. 16, 552–561. https://doi.org/10.1038/nn.3359

Wang, S., Tang, N.H., Lara-Gonzalez, P., Zhao, Z., Cheerambathur, D.K., Prevo, B., Chisholm, A.D., Desai, A., Oegema, K., 2017. A toolkit for GFP-mediated tissue-specific protein degradation in C. elegans . Development 144, 2694–2701. https://doi.org/10.1242/dev.150094

Wickham, H., 2016. ggplot2: Elegant Graphics for Data Analysis. Springer-Verlag, New York.

Willemsen, M.H., Nijhof, B., Fenckova, M., Nillesen, W.M., Bongers, E.M.H.F., Castells-Nobau, A., Asztalos, L., Viragh, E., van Bon, B.W.M., Tezel, E., Veltman, J.A., Brunner, H.G., de Vries, B.B.A., de Ligt, J., Yntema, H.G., van Bokhoven, H., Isidor, B., Le Caignec, C., Lorino, E., Asztalos, Z., Koolen, D.A., Vissers, L.E.L.M., Schenck, A., Kleefstra, T., 2013. *GATAD2B* loss-of-function mutations cause a recognisable syndrome with intellectual disability and are associated with learning deficits and synaptic undergrowth in *Drosophila* J. Med. Genet. 50, 507 LP – 514. https://doi.org/10.1136/jmedgenet-2012-101490

Williams, L.E., Blackford, J.U., Luksik, A., Gauthier, I., Heckers, S., 2013. Reduced habituation in patients with schizophrenia. Schizophr. Res. 151, 124–132. https://doi.org/10.1016/j.schres.2013.10.017

Willsey, A.J., Sanders, S.J., Li, M., Dong, S., Tebbenkamp, A.T., Muhle, R.A., Reilly, S.K., Lin, L., Fertuzinhos, S., Miller, J.A., Murtha, M.T., Bichsel, C., Niu, W., Cotney, J., Ercan-Sencicek, A.G., Gockley, J., Gupta, A.R., Han, W., He, X., Hoffman, E.J., Klei, L., Lei, J., Liu, W., Liu, L., Lu, C., Xu, X., Zhu, Y., Mane, S.M., Lein, E.S., Wei, L., Noonan, J.P., Roeder, K., Devlin, B., Sestan, N., State, M.W., 2013. Coexpression Networks Implicate Human Midfetal Deep Cortical Projection Neurons in the Pathogenesis of Autism. Cell 155, 997–1007. https://doi.org/10.1016/j.cell.2013.10.020

Xiang, M., Zhou, L., Macke, J.P., Yoshioka, T., Hendry, S.H., Eddy, R.L., Shows, T.B., Nathans, J., 1995. The Brn-3 family of POU-domain factors: primary structure, binding specificity, and expression in subsets of retinal ganglion cells and somatosensory neurons. J. Neurosci. 15, 4762–85.

Zeier, Z., Kumar, A., Bodhinathan, K., Feller, J.A., Foster, T.C., Bloom, D.C., 2009. Fragile X mental retardation protein replacement restores hippocampal synaptic function in a mouse model of fragile X syndrome. Gene Ther. 16, 1122–1129. https://doi.org/10.1038/gt.2009.83

Zhang, L., Ward, J.D., Cheng, Z., Dernburg, A.F., 2015. The auxin-inducible degradation (AID) system enables versatile conditional protein depletion in C. elegans. Development 142, 4374–4384. https://doi.org/10.1242/dev.129635

Zou, M., Li, S., Klein, W.H., Xiang, M., 2012. Brn3a/Pou4f1 regulates dorsal root ganglion sensory neuron specification and axonal projection into the spinal cord. Dev. Biol. 364, 114–127. https://doi.org/10.1016/j.ydbio.2012.01.021

